# LKB1 is the gatekeeper of carotid body chemo-sensing and the hypoxic ventilatory response

**DOI:** 10.1101/2022.05.24.493275

**Authors:** Sandy MacMillan, Andrew P. Holmes, Mark L. Dallas, Amira D. Mahmoud, Michael J. Shipston, the late Chris Peers, D. Grahame Hardie, Prem Kumar, A. Mark Evans

## Abstract

The hypoxic ventilatory response (HVR) is critical to breathing and thus oxygen supply to the body and is primarily mediated by the carotid bodies. Here we reveal that carotid body afferent discharge during hypoxia and hypercapnia is determined by the expression of Liver Kinase B1 (LKB1), the principal kinase that activates the AMP-activated protein kinase (AMPK) during metabolic stresses. Conversely, conditional deletion in catecholaminergic cells of AMPK had no effect on carotid body responses to hypoxia or hypercapnia. By contrast, the HVR was attenuated by LKB1 and AMPK deletion. However, in LKB1 knockouts hypoxia evoked hypoventilation, apnoea and Cheyne-Stokes-like breathing, while only hypoventilation and apnoea were observed after AMPK deletion. We therefore identify LKB1 as an essential regulator of carotid body chemosensing and uncover a divergence in dependency on LKB1 and AMPK between the carotid body on one hand and the HVR on the other.

## Introduction

The hypoxic ventilatory response (HVR) delivers compensatory increases in ventilatory drive where there are deficiencies in oxygen availability. The HVR is initiated by increases in afferent fibre discharge from the carotid bodies, the primary peripheral arterial chemoreceptors of mammals that are located within the bifurcations of the carotid artery, ideally situated to monitor blood flow to the brain. Within the carotid body falls in arterial *P*O_2_ (and increases in arterial *P*CO_2_) are sensed directly by carotid body type I cells, where consequent depolarisation elicits exocytotic release of ATP that mediates increases in chemoafferent discharge to the respiratory central pattern generators in the brainstem^1–3^.

We recently showed that in addition to regulating metabolic homeostasis in a cell autonomous manner^4^ the AMP-activated protein kinase (AMPK) facilitates the HVR and thus oxygen and energy (ATP) supply to the whole body^5^. In doing so, we demonstrated that AMPK acts not at the level of the carotid bodies as one would predict but downstream at the brainstem. Briefly, conditional deletion of AMPK in catecholaminergic neurons of mice precipitates hypoventilation and apnoea during poikilocapnic hypoxia^5^, that resembles central apnoea of prematurity^6, 7^ and central sleep apnoea^8^ in neonate and adult humans, respectively. That said, the HVR of these mice is most reminiscent of the HVR observed in premature infants where the hypercapnic hypoxic ventilatory response is similarly conserved^9^. Given such face validity that resembles symptoms in patients, it is important that we identify the mechanism(s) by which AMPK is regulated in the context of the HVR.

The principal pathway of AMPK activation by metabolic stresses is through direct phosphorylation by Liver Kinase B1 (LKB1), which exists in a complex with regulatory proteins STRAD and MO25^10–12^. LKB1 also regulates by direct phosphorylation eleven of the twelve AMPK-related kinases^13^, but in each case this is insensitive to metabolic stresses^14^. Only AMPK is coupled to LKB1 through changes in the cellular AMP:ATP and ADP:ATP ratios^15^, that may be triggered through inhibition of mitochondrial oxidative phosphorylation during hypoxia, as is the HVR^16, 17^. Binding of AMP to the AMPK-γ subunit increases activity 10-fold by allosteric activation alone, while binding of AMP or ADP triggers increases in phosphorylation of Thr172 on the α subunit by LKB1 (conferring up to 100-fold further activation) and at the same time reduces Thr172 dephosphorylation^18^. However, alternative AMP-independent mechanisms of AMPK activation have been identified: (i) calcium-dependent Thr172 phosphorylation by the calmodulin-dependent protein kinase CaMKK2^19^; (ii) long chain fatty acyl-CoA binding to the Allosteric Drug & Metabolite (ADaM) site on the α subunit^20^; (iii) glucose-deprivation^21^.

Clearly, the most likely path to AMPK activation during hypoxia would be through increases in the AM(D)P:ATP ratio and thus LKB1-dependent phosphorylation. However, the fact that AMPK facilitates the HVR within regions of the brainstem that receive carotid body afferent input responses^5^, rather than at the level of the carotid bodies^1–3^, also suggests a role for the alternative CaMKK2 pathway, which has been proposed to contribute to energy balance regulation by hypothalamic networks^22^.

We set out to examine the mechanism by which AMPK is regulated in the context of the HVR. To this end we employed a three-point assay to assess the relationship between the level of LKB1 expression, carotid body activation during hypoxia and the HVR, by utilising a mouse line which exhibits ∼90% global hypomorphic expression of the gene that encodes LKB1 (*Stk11*, hereafter referred to as *Lkb1*)^23, 24^ and a conditional homozygous LKB1 knockout mouse line derived from this in which 100% *Lkb1* deletion is triggered by Cre expression via the tyrosine hydroxylase (TH) promoter, which restricts *Lkb1* deletion to catecholaminergic cells including therein carotid body type I cells. We compared outcomes to those observed in mice where CaMKK2 had been deleted globally. The present investigation not only demonstrates that LKB1, but not CaMKK2, is required for the HVR, but also reveals that the level of LKB1 expression serves an essential role in establishing carotid body function and chemosensitivity. In short, we uncover a divergence in dependency on LKB1 and AMPK between the carotid body on the one hand and the HVR on the other. Adding to this we show that LKB1, but not AMPK, deficiency within catecholaminergic cells precipitates Cheyne-Stokes-like breathing patterns during hypoxia, which are associated with heart failure but of unknown aetiology^25^.

## Results

### LKB1 deficiency augments while homozygous *Lkb1* gene deletion ablates type I cell activation in response to hypoxia

We confirmed that Cre expression and deletion of the gene encoding LKB1 was targeted to catecholaminergic cells by two means. Firstly, these mice were crossed with a mouse line expressing the Cre-inducible reporter gene Rosa (tdTomato), the expression of which in type I cells was assessed by confocal imaging of acutely isolated and fixed sections of tissue comprising the superior cervical ganglion, carotid artery and carotid body (Fig 1a; note, tyrosine hydroxylase is expressed by carotid body type I cells, endothelial cells and sympathetic neurons). Then, the absence of LKB1 expression was confirmed in acutely isolated carotid body type I cells by single cell end point RT-PCR (Fig 1b-c **and Supplementary Fig 1**); consistent with outcomes for other organs including the brain^23, 24^ LKB1 expression from homozygous *Lkb1* floxed mice (Ct = 31.147 ± 0.098, n = 3) was lower than for TH-Cre mice (Ct = 25.139 ± 0.006, n = 3) in whole carotid bodies, although it should be noted that this multi-cellular organ is not representative of pure type I cells.

**Figure 1.**
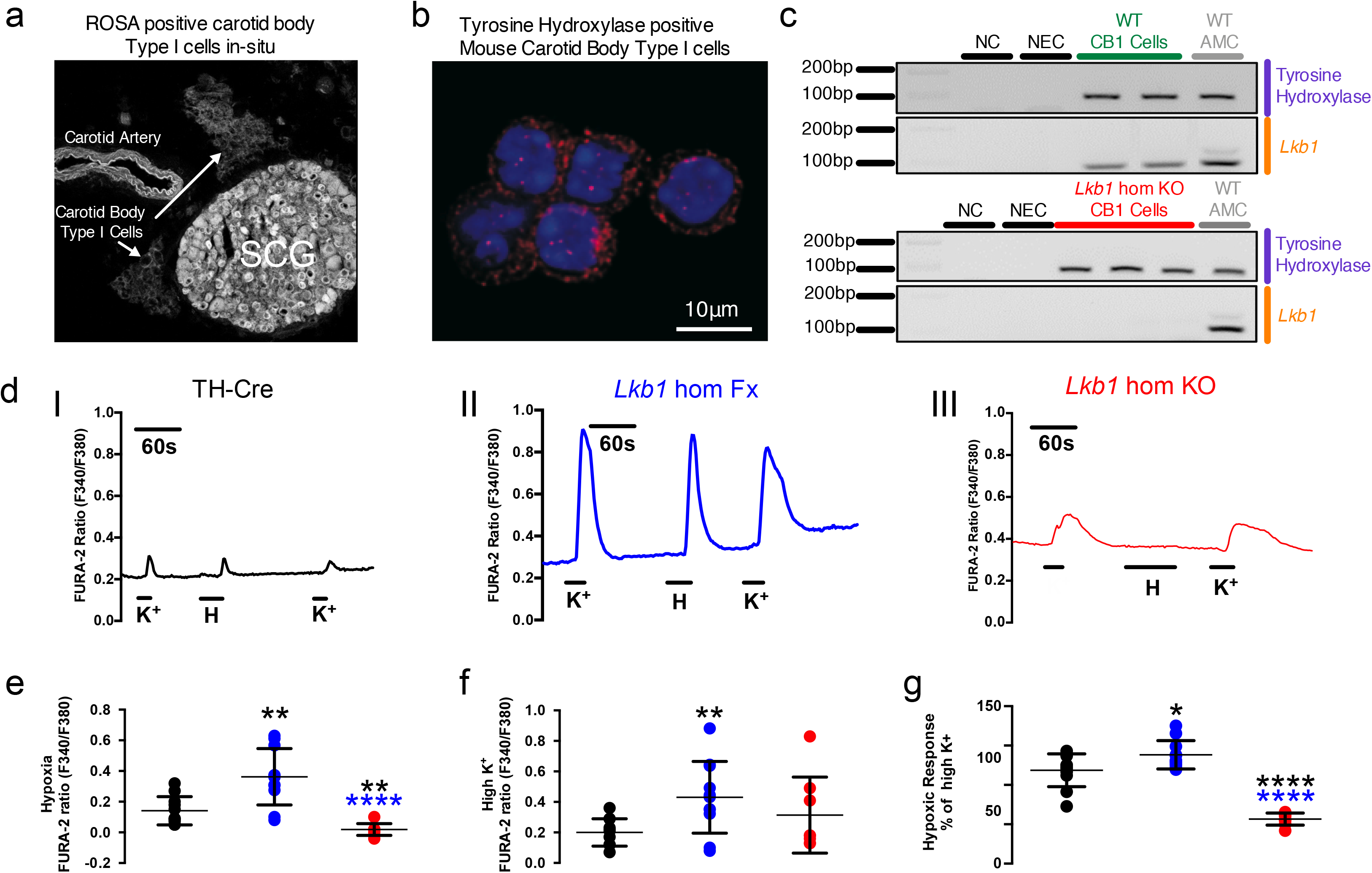
Conditional deletion of *Lkb1* in carotid body type I cells blocks hypoxia-evoked calcium transients. a, Confocal image shows tdTomato (excitation 555 nm, emission 582 nm) positive type I cells in-situ within a section of tissue comprising the carotid body, superior cervical ganglion (SCG) and carotid artery; note endothelial cells and sympathetic neurons express tyrosine hydroxylase (TH). b, Acutely isolated carotid body type I cells stained for TH and DAPI. c, Single cell end-point RT-PCR amplicons for TH and *Lkb1* from acutely isolated adrenal medullary chromaffin cells (WT AMCs) and carotid body type I cells of wild type (WT CB1 cells) and conditional *Lkb1* knockout mice (*Lkb1* hom KO CB1 cells); NC = negative control (cell aspirant but no reverse transcriptase added); NEC = negative extracellular control (aspirant of extracellular medium). d, Exemplar records show calcium transients evoked by 50 mM potassium and hypoxia (mean±SEM *P*O_2_ = 20.19±1.73mmHg, ∼2% O_2_; n = 10) in type I cells isolated from (I) TH-Cre (black; n = 8 different carotid body type I cells), (II) homozygous *Lkb1* homozygous floxed (*Lkb1* hom Fx, blue; n = 11 different carotid body type I cells) and (III) conditional homozygous *Lkb1* hom KO (red, n= 8 different carotid body type I cells) mice. e-g, Dot plots show mean±SEM F340/F380 ratios for calcium transients evoked by (e) 50 mM potassium, (f) hypoxia, while (g) shows the hypoxic response expressed as a ratio of the response to 50mM potassium. *=p<0.05, **=p<0.01, ****=p<0.0001. Replicates taken from ≥3 different mice.

We next examined the impact of LKB1 deletion on carotid body type I cell function. To this end we employed TH-Cre mice as the control group, because these mice were used to deliver *Lkb1* deletion in catecholaminergic cells and there was no significant difference between the hypoxic ventilatory response of these mice when compared to the background strain (C57/BL6; **Supplementary Fig 2**). Changes in intracellular calcium concentration within isolated type I cells were assessed as an index of their activation by hypoxia, using the ratiometric calcium indicator Fura-2. Hypoxia (mean±SEM *P*O_2_ = 20.19±1.73mmHg, ∼2% O_2_ (n=10); from normoxia ∼150mmHg, ∼21% O_2_) induced a robust increase in Fura-2 fluorescence ratio in type I cells from TH-Cre mice (n = 8), which was equivalent to that resulting from voltage-gated calcium influx triggered by membrane depolarisation in response to extracellular application of 50mM potassium chloride (Fig 1di **and** e-g**, Supplementary Fig 3**). Surprisingly, outcomes for type I cells from *Lkb1* floxed mice and *Lkb1* knockouts were not only different but opposite. Potassium-evoked calcium transients were markedly augmented in *Lkb1* floxed mice (n = 11; Fig 1dii **and** e-g**, Supplementary Fig 3**), which are hypomorphic with ∼90% lower expression of LKB1 globally when compared to wild type controls^23^. In marked contrast, hypoxia failed to increase intracellular calcium in type I cells from homozygous *Lkb1* knockouts (n = 8), where calcium transients evoked by 50mM potassium chloride were equivalent to controls (Fig 1diii **and** e-g).

### LKB1 deficiency attenuates while homozygous *Lkb1* deletion virtually abolishes increases in carotid body afferent discharge during hypoxia and hypercapnia

Extracellular recordings of single unit activity from the carotid sinus nerve, showed that increases in afferent discharge frequency during hypoxia were attenuated in carotid bodies from *Lkb1* floxed mice, and virtually abolished in carotid bodies from *Lkb1* knockouts (Fig 2a-b).

**Figure 2.**
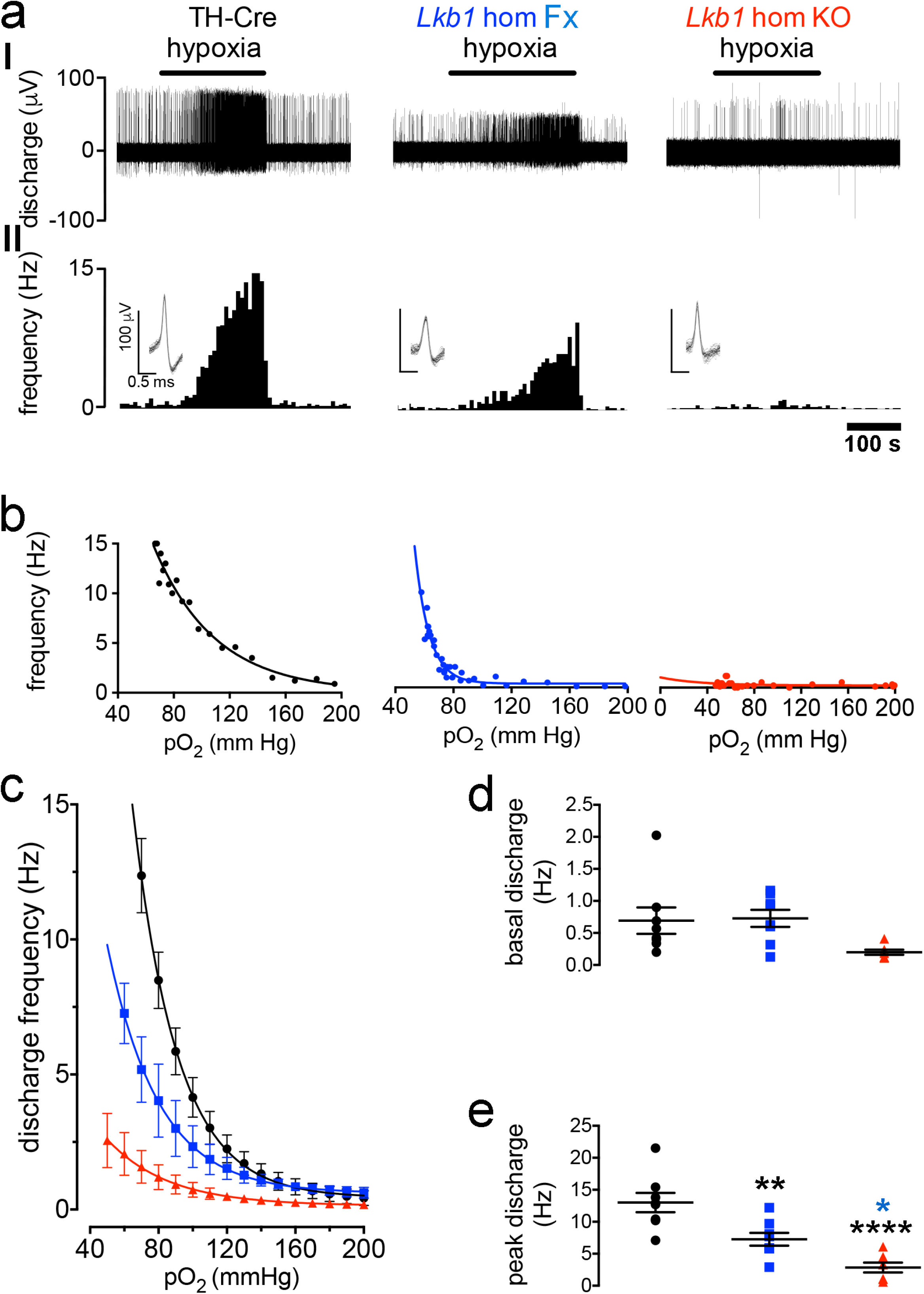
Conditional deletion of *Lkb1* in carotid body type I cells attenuates basal and hypoxia-evoked afferent discharge from the carotid body in-vitro. a shows (I) extracellular recordings of chemoafferent discharge versus time during normoxia and hypoxia and (II) frequency-time histograms (*inset:* single fibre discriminations) for carotid bodies from control (TH-Cre, black), *Lkb1* homozygous floxed (*Lkb1* hom Fx, blue, middle panels), and conditional *Lkb1* homozygous knockout mice(*Lkb1* hom KO, red). b, Exemplar frequency-*P*O_2_ response curves for records shown in (a). c, Compares mean±SEM for frequency-*P*O_2_ response curves for TH-Cre (n = 8 different carotid bodies), homozygous *Lkb1* floxed (n = 8 different carotid bodies) and conditional homozygous *Lkb1* knockout (n = 7 different carotid bodies) mice. Dot plots show mean±SEM for (d) basal single fibre discharge frequency and (e) peak single fibre discharge frequency during hypoxia. * =p<0.05, ** =p<0.01, ****=p< 0.0001.

During normoxia, basal afferent fibre discharge frequency from in-vitro carotid bodies of controls (TH-Cre, n=8) was similar to that of homozygous *Lkb1* floxed mice (p = 0.38 by ANOVA; p = 0.17 by Student’s t test; n = 8; Fig 2d) that exhibit ∼90% global reductions in LKB1 expression^23^. By contrast, mean basal afferent discharge from carotid bodies of homozygous *Lkb1* knockouts (n=7) was reduced by approximately 70% (Fig 2d) which reached significance by Student’s t test (p<0.05 versus TH-Cre) but not by ANOVA (p = 0.09 versus TH-Cre and p = 0.06 vs *Lkb1* floxed).

Reductions in superfusate *P*O_2_ evoked exponential increases in afferent discharge from carotid bodies of controls and *Lkb1* floxed mice. Intriguingly, however, during hypoxia (*P*O_2_ ≤75mmHg) peak discharge frequencies of *Lkb1* floxed mice (∼90% loss of LKB1 expression) were attenuated by ∼50% relative to controls (TH-Cre; p<0.01; Fig 2c and e). Furthermore, the *P*O_2_ required to reach a frequency of 5 Hz was lower in the *Lkb1* floxed mice (70±7 mmHg) compared to TH-Cre controls (96±4 mmHg, p<0.01) (**Supplementary Fig 3)**, indicative of a delay / lower *P*O_2_ threshold for response initiation. Exponential rate constants were consistent between TH-Cre and homozygous *Lkb1* floxed mice (**Supplementary Fig 3**). By contrast, reductions in superfusate *P*O_2_ evoked little or no increase in afferent discharge from carotid bodies of homozygous *Lkb1* knockouts (Fig 2c and e; p<0.0001 versus TH-Cre and p<0.05 versus *Lkb1* floxed), consistent with the fact that homozygous *Lkb1* deletion abolished type I cell activation during hypoxia. One can only assume that the ∼50% reduction in hypoxia-evoked afferent discharge in carotid bodies isolated from *Lkb1* floxed mice was due to their ∼90% deficiency in LKB1 expression^23^, despite the fact that hypoxia-evoked calcium transients in isolated type I cells from these mice were augmented relative to controls (see Fig 1).

Our original assumption had been that LKB1 would primarily function to couple reductions in mitochondrial ATP supply to carotid body type I cell activation and thus carotid body afferent discharge during hypoxia. It was a surprise to find, therefore, that *Lkb1* deletion also attenuated carotid body activation during hypercapnia (Fig. 3a), given that carotid body responses to hypercapnia are generally presumed to be triggered by ion channel mechanisms independent of mitochondrial metabolism^26–29^, although earlier studies had provided pharmacological evidence that pointed to a partial dependence on mitochondria (see Discussion for further details)^30^. CO_2_-sensitivity (which is linear between 40 and 80 mmHg) was largely preserved in the *Lkb1* floxed mice (n=6, p = 0.38 versus TH-Cre) but was virtually abolished in carotid bodies from homozygous *Lkb1* knockouts (n=4) when compared to TH-Cre (p<0.01 versus TH-Cre; p<0.06 versus *Lkb1 floxed*; n = 7; Fig 3b and c).

**Figure 3.**
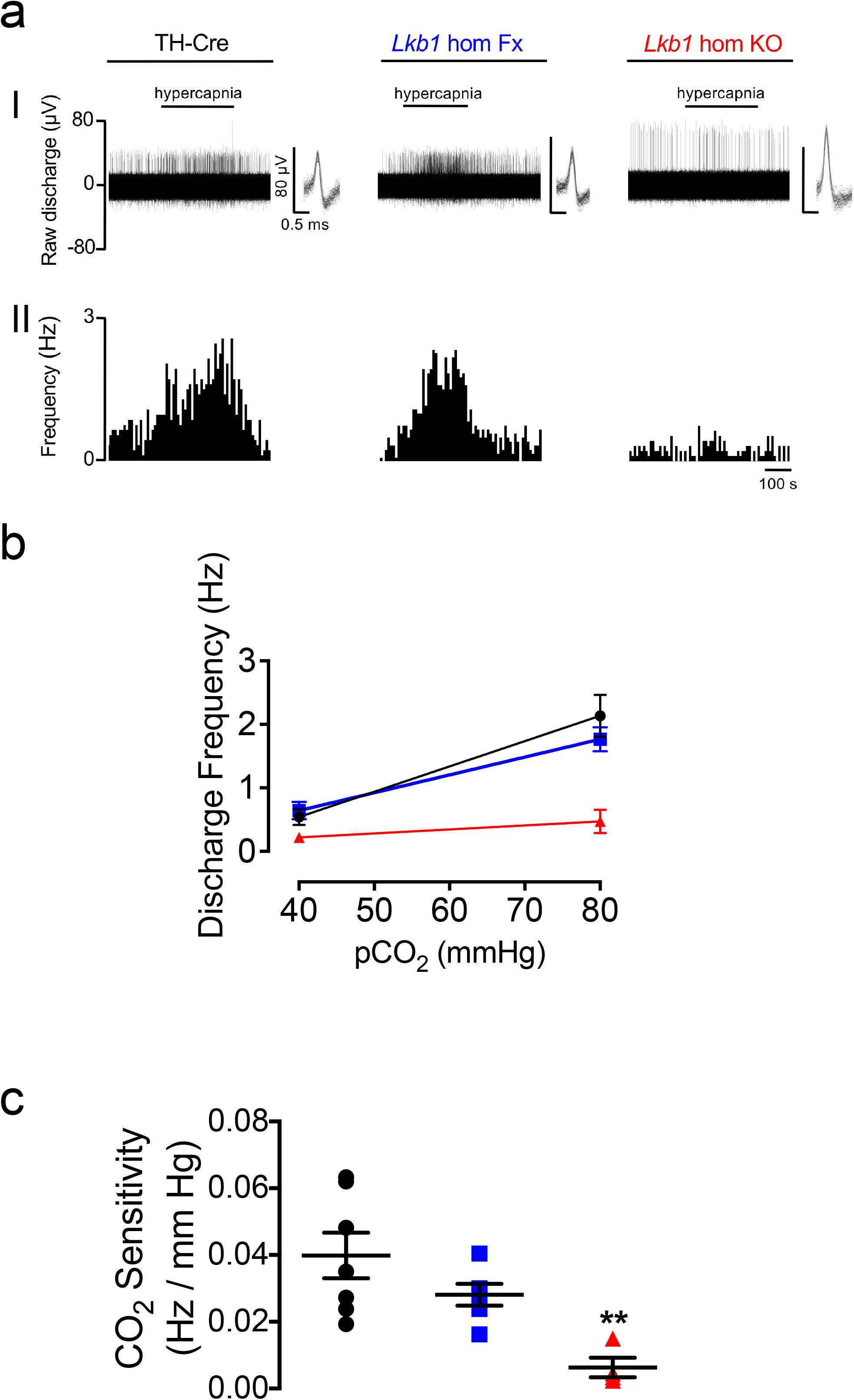
Conditional deletion of *Lkb1* in carotid body type I cells attenuates basal and hypercapnia-evoked afferent discharge from the carotid body in-vitro. a, shows (I) extracellular recordings of chemoafferent discharge versus time during normoxia/normocapnia and hypercapnia and (II) frequency-time histograms for carotid bodies from control (TH-Cre, black; n = 7 different carotid bodies), homozygous *Lkb1* floxed (*Lkb1* hom Fx, blue; n = 6 different carotid bodies) and conditional *Lkb1* homozygous knockout (*Lkb1* hom KO, red; n = 4 different carotid bodies) mice (*inset:* single fibre discriminations). b shows mean±SEM for chemoafferent discharge versus *P*CO_2_. c, Dot plots show mean±SEM for CO_2_ sensitivity for TH-Cre (black), *Lkb1* hom Fx (blue) and *Lkb1* hom KO (blue). **=p< 0.01.

In stark contrast, and consistent with our preliminary findings^5^, extracellular recordings of single unit activity from the carotid sinus nerve, showed that increases carotid body afferent responses to hypoxia in the *AMPKα1+α2* homozygous knockouts (n=8) remained comparable to TH-Cre (n=8) and *AMPKα1+α2* homozygous floxed (n = 10) controls (Fig 4a-c**, Supplementary Fig 4;** for single cell PCR see^5^). Basal discharge and peak responses to hypoxia were similar between the three groups (Fig 4a-c**, Supplementary Fig 4**) although a significant reduction in basal discharge was detected for the *AMPKα1+α2* homozygous knockouts when compared to *AMPKα1+α2* homozygous floxed mice by Student’s t test (p<0.05) but not ANOVA. The *P*O_2_ required to reach a frequency of 5Hz was reduced in the *AMPKα1+α2* homozygous floxed mice (73±5 mmHg, p<0.01) and showed a similar trend in the *AMPKα1+α2* homozygous knockouts (80±5 mmHg, p = 0.08) compared to TH-Cre controls (96±4 mmHg; **Supplementary Fig 4**). There was, however, no difference in the *P*O_2_ required to reach a frequency of 5Hz between *AMPKα1+α2* homozygous floxed mice and *AMPKα1+α2* homozygous knockouts. Exponential rate constants were consistent between the three groups (**Supplementary Fig 4)**. Moreover, carotid body responses to hypercapnia and CO_2_-sensitivity remained unaltered in *AMPKα1+α2* homozygous knockouts (n=9) compared to TH-Cre (n=7) and *AMPKα1+α2* homozygous floxed mice (n=9; Fig 4d-f**, Supplementary Fig 4**).

**Figure 4.**
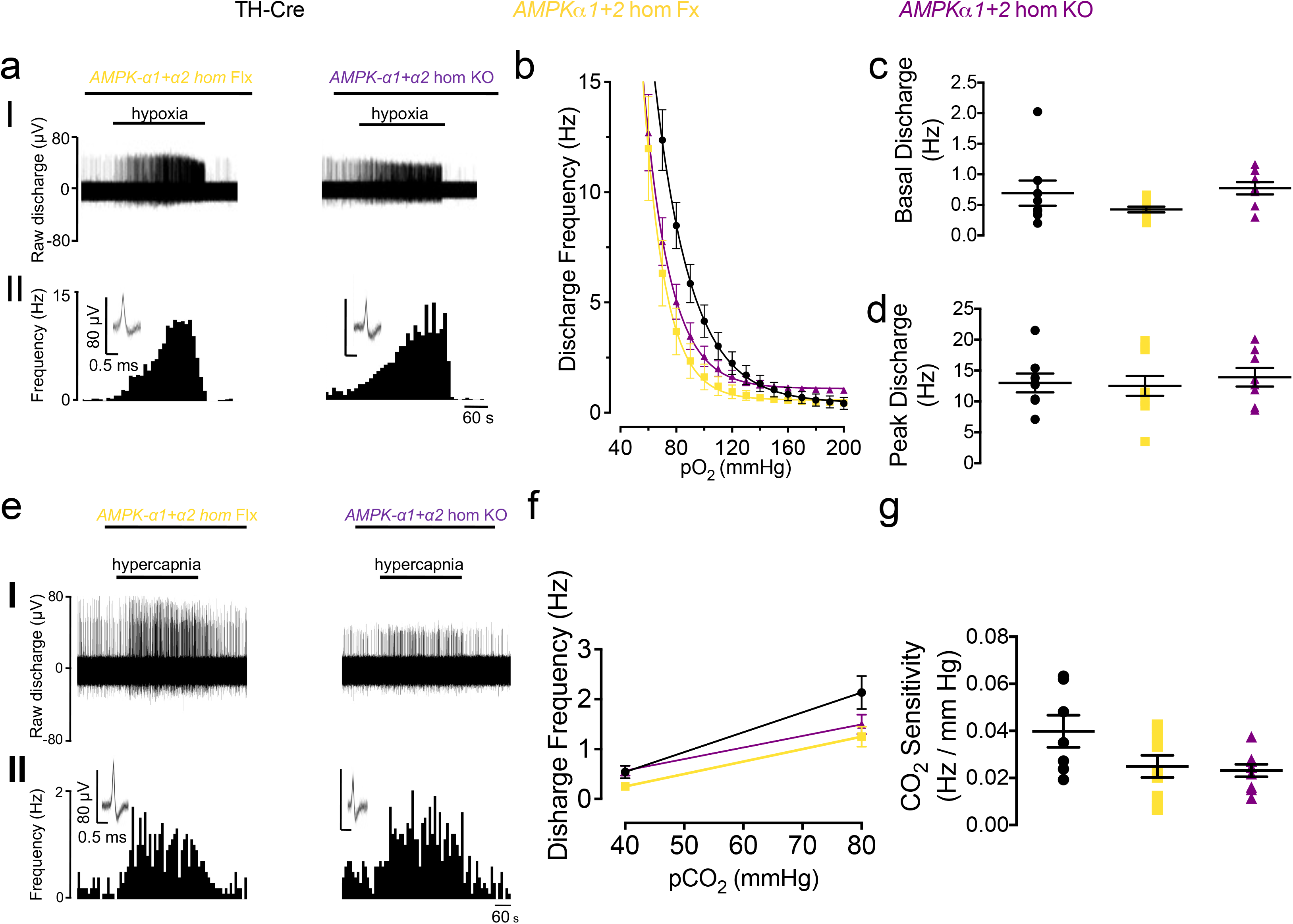
Conditional deletion of *AMPK-α1+*α2 in carotid body type I cells has no effect on hypoxia-evoked or hypercapnia-evoked afferent discharge from the carotid body in-vitro. a, Shows (I) extracellular recordings of chemoafferent discharge versus time during normoxia and hypoxia and (II) frequency-time histograms (*inset:* single fibre discriminations) for carotid bodies from controls (TH-Cre, black; n = 8 different carotid bodies), *AMPKα1+α2* homozygous floxed *(AMPKα1+α2* hom Fx, beige; n = 9 different carotid bodies) and conditional homozygous *AMPKα1+α2* knockout mice (*AMPKα1+α2* hom KO, purple; n = 9 different carotid bodies). b, Means±SEM for frequency-*P*O_2_ response curves for homozygous *AMPKα1+α2* hom Fx and *AMPKα1+α2* hom KO. c-d, Dot plots show mean±SEM for (c) basal single fibre discharge frequency and (d) peak single fibre discharge frequency during hypoxia. e, as for (a) but in response to hypercapnia. f, Means±SEM for frequency-*PC*O_2_ relationship. g, Dot plot shows mean±SEM for CO_2_ sensitivity.

Taken together, these findings suggest that LKB1 establishes, independent of AMPK, carotid body sensitivity to hypoxia and hypercapnia through a related mechanism, the set point of which can be adjusted by changes in LKB1 expression. By contrast, within type I cells AMPK may support an inhibitory input on basal afferent discharge during normoxia.

### The HVR is attenuated in mice with LKB1 deficiency but remains unaffected following *Camkk2* deletion

Under normoxia there was no difference between controls and either *Lkb1*, *Camkk2* or, as previously shown^5^, dual *AMPKα1+α2* knockouts with respect to breathing frequency, tidal volume, minute ventilation, blood gases, blood pH or core body temperature (**Supplementary Fig 5 and Table 1**). Moreover, the metabolic status of *Lkb1* floxed mice is normal^23^. Nevertheless, profound genotype-specific differences were observed with respect to the ventilatory responses during hypoxia, and to a lesser extent during hypercapnia (**Supplementary Movies 1-4**).

During exposures to poikilocapnic hypoxia the peak of the initial “Augmenting Phase” of the HVR (∼30s) remained unaffected in *Lkb1* floxed mice (12% O_2_, n = 14; 8% O_2_, n = 16) that harbour ∼90% global reductions in LKB1 expression^23^ when compared to TH-Cre (n=25). By contrast, the subsequent “Sustained Phase” (2-5 min) of the HVR was attenuated during severe poikilocapnic hypoxia (8% O_2_; p<0.0001 relative to TH-Cre; n=37) but not during moderate (12% O_2_; n = 25; 8% O_2_, n = 37) poikilocapnic hypoxia (Fig 5a-b); i.e., these mice exhibited delayed hypoventilation during severe hypoxia.

**Figure 5.**
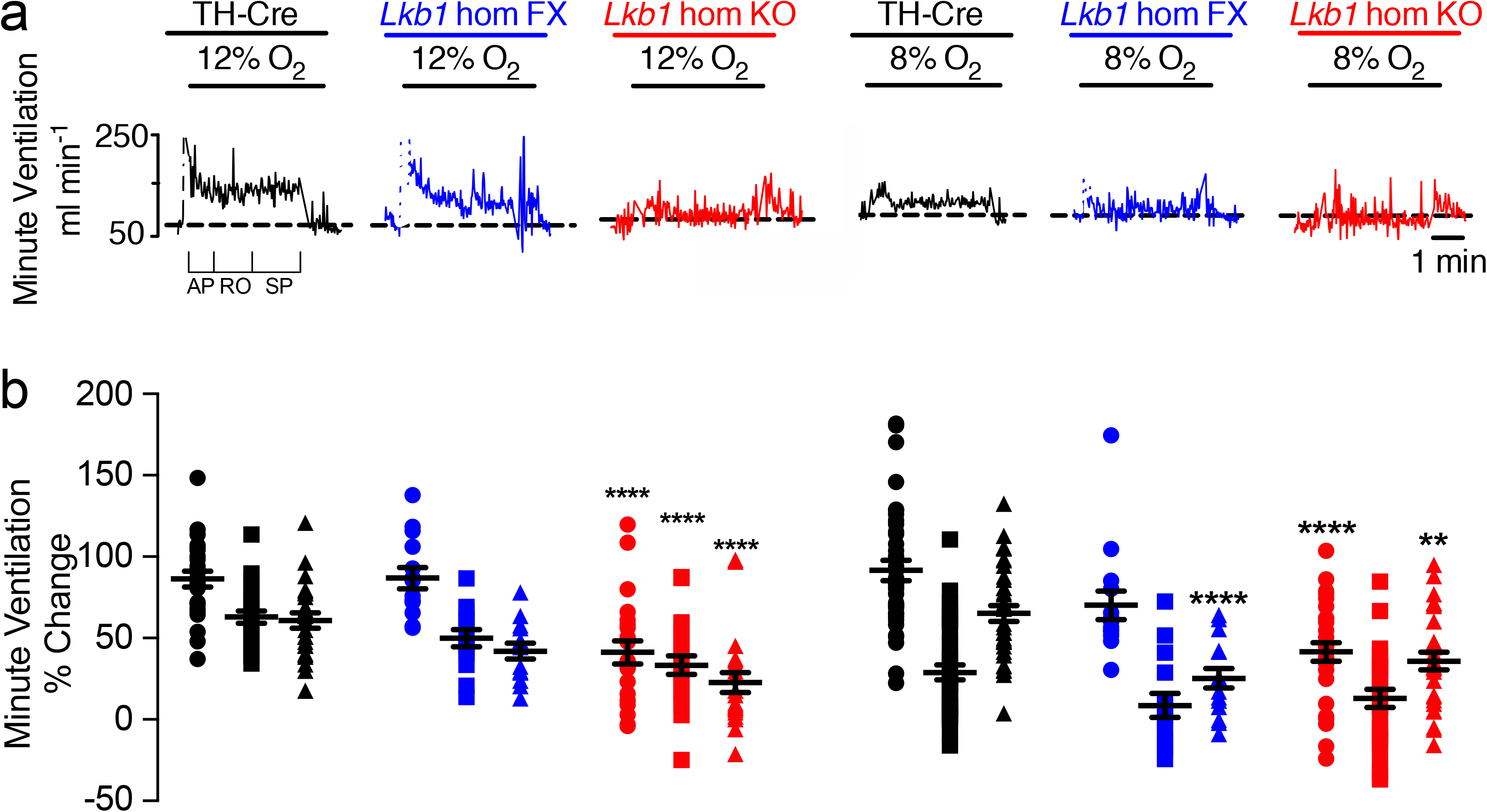
Mice hypomorphic for LKB1 exhibit an attenuated hypoxic ventilatory response measured by unrestrained plethysmography. a, Example records of minute ventilation versus time. b, Dot plots of mean±SEM for % change in minute ventilation at the peak of the Augmenting Phase (AP, ∼30s), after Roll Off (RO, ∼100s) and during the plateau of the Sustained Phase (SP, ∼300s) of the ventilatory response to 12% and 8% O_2_ for TH-Cre (black; 12% O_2_, n = 25 independent experiments; 8% O_2_, n = 37 independent experiments), *Lkb1* homozygous floxed (*Lkb1* hom Fx, blue; 12% O_2_, n = 14 independent experiments; 8% O_2_, n = 15 independent experiments) and conditional *Lkb1* homozygous knockout mice (*Lkb1* hom KO, red; n = 22 independent experiments; 8% O_2_, n = 30 independent experiments). **=p<0.01; ****=p<0.0001.

The effect of conditional *Lkb1* deletion in catecholaminergic cells was more severe (Fig 5a-b). The HVR was suppressed throughout 5min exposures to either mild (Fig 5b; 12% O_2_, n = 14) or severe hypoxia (Fig 5b; 8% O_2_, n = 15), and in a manner related to the severity of hypoxia. In marked contrast to the outcomes for mice with hypomorphic expression of LKB1, complete *Lkb1* deletion markedly attenuated the peak change in minute ventilation of the initial “Augmenting Phase” of the HVR during mild and severe hypoxia (at ∼30s; p<0.0001 compared to TH-Cre), which is primarily driven by carotid body afferent input responses ^1, 31, 32^. Following subsequent ventilatory depression (Roll Off, ∼100s) the HVR was attenuated during mild hypoxia (p<0.0001, compared to TH-Cre), but this did not reach significance during severe hypoxia. However, the latter Sustained Phase of the HVR (2-5min) was markedly attenuated during mild (p<0.0001, compared to TH-Cre) and severe hypoxia (p<0.01, compared to TH-Cre). Note, the 0.05% CO_2_ used here was probably insufficient to prevent respiratory alkalosis which may have impacted on ventilatory reflexes during the latter phases of the sustained hypoxic stimulus ^33^, in wild type mice in particular. Therefore, we may have underestimated the degree to which *Lkb1* deletion inhibits the HVR.

By contrast to the effects of *Lkb1* deletion, global deletion of *Camkk2* (n = 10) had no discernable effect on the HVR (**Supplementary Fig 6**), ruling out a prominent role for CaMKK2 in facilitating the acute HVR alone or through AMPK activation^19^.

More detailed analysis in *Lkb1* floxed mice identified attenuation of increases in breathing frequency at all time points during exposure to severe (8% O_2_, n = 22) but not mild hypoxia (12% O_2_, n = 15), including therein the Augmenting Phase (p<0.05 relative to TH-Cre), Roll Off (p<0.0001 relative to TH-Cre) and the Sustained Phase (p<0.0001 relative to TH-Cre). Increases in breathing frequency during hypoxia were yet more markedly attenuated by homozygous *Lkb1* deletion throughout exposures to both mild and severe hypoxia (p<0.0001 relative to TH-Cre; Fig 6a), and in a manner proportional to the severity of hypoxia. By contrast no attenuation of increases in tidal volume was observed for either *Lkb1* floxed mice or *Lkb1* knockouts during mild or severe hypoxia (Figure 6b). In fact, during severe hypoxia *Lkb1* deletion in catecholaminergic cells, but not hypomorphic expression of LKB1, appeared to augment increases in tidal volume during Roll Off and the Sustained Phase (p<0.01 relative to TH-Cre), but not during the initial Augmenting Phase.

**Figure 6.**
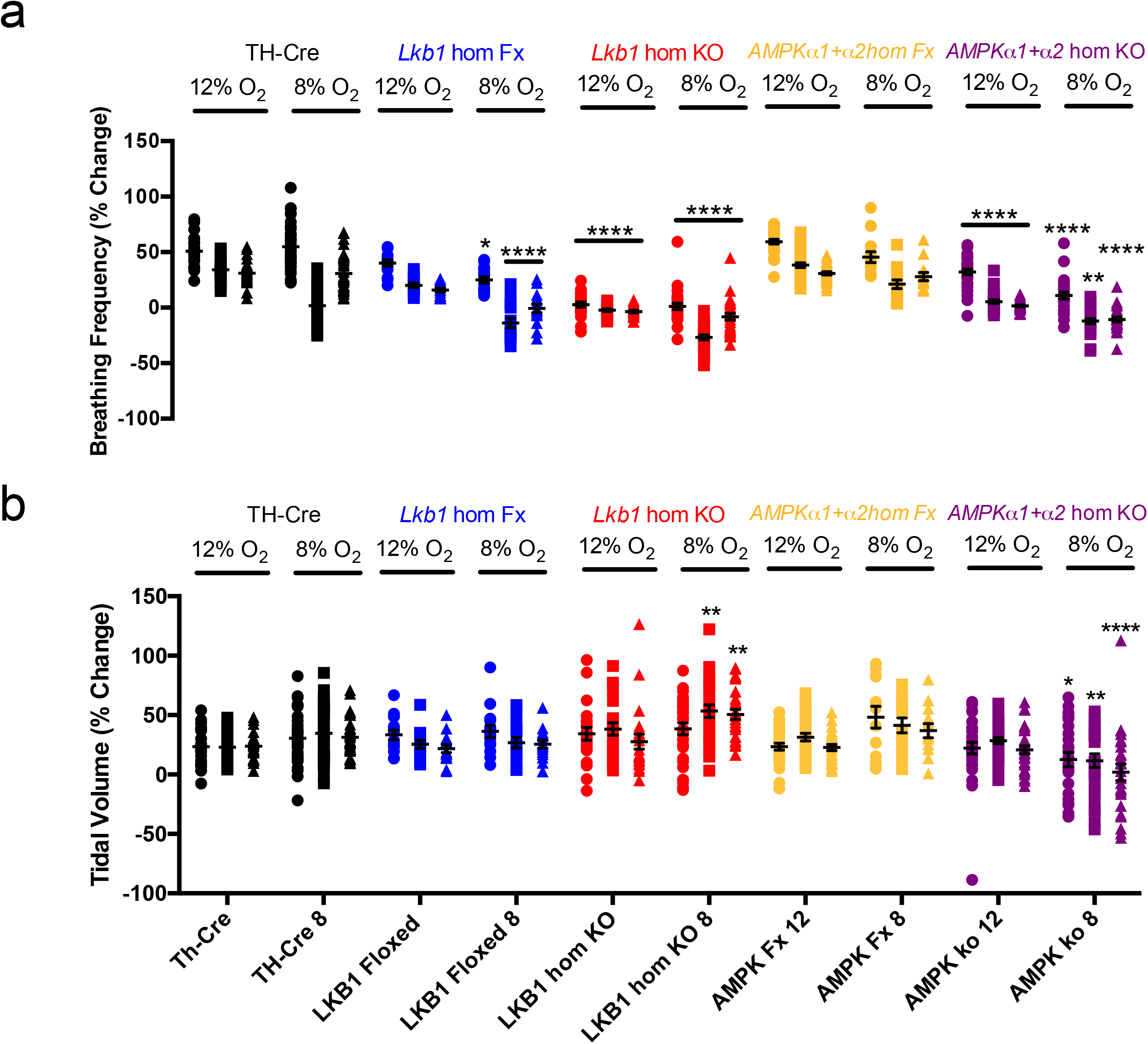
Conditional deletion of *Lkb1* and AMPK in tyrosine hydroxylase expressing cells attenuates increases in breathing frequency during hypoxia but only *Lkb1* deletion augments increases in tidal volume during severe hypoxia. Dot plots of mean±SEM for changes in (a) breathing frequency and (b) tidal volume at the peak of the Augmenting Phase (∼30s), at ∼100s following Roll Off and during the plateau of the Sustained Phase (∼300s) of the ventilatory response to mild (12% O_2_) and severe (8% O_2_) hypoxia for TH-Cre (black; 12% O_2_, n = 25 independent experiments; 8% O_2_, n = 37 independent experiments), *Lkb1* homozygous floxed (*Lkb1* hom Fx, blue; 12% O_2_, n = 14 independent experiments; 8% O_2_, n = 15 independent experiments) that are ∼90% hypomorphic for LKB1 and conditional *Lkb1* homozygous knockout mice (*Lkb1* hom KO, red; 12% O_2,_ n = 22 independent experiments; 8% O_2_, n = 30 independent experiments). These data are also compared with outcomes for *AMPKα1*+*α2* homozygous floxed mice (*AMPKα1*+*α2* hom Fx, beige, 12% O_2_ n = 30 independent experiments; 8% O_2_, n = 13 independent experiments) and conditional *AMPKα1*+*α2* homozygous knockout mice (*AMPKα1*+*α2* hom KO, purple, 12% O_2_ n = 30 independent experiments; 8% O_2_, n = 26 independent experiments). *=p<0.05, **=p<0.01, ****=p<0.0001 compared to TH-Cre.

Conditional deletion of *AMPKα1+α2* in catecholaminergic cells also attenuated increases in breathing frequency, but not tidal volume, when these mice were exposed to mild hypoxia (12% O_2;_ Fig 6a-b; n = 30) including therein the Augmenting Phase (p<0.0001 relative to TH-Cre), Roll Off (p<0.0001 for 12% O_2_; p<0.001 for 8% O_2_; relative to TH-Cre) and the Sustained Phase (p<0.0001 relative to TH-Cre). By contrast to the impact of *Lkb1* deletion, however, *AMPKα1+α2* deletion attenuated increases in tidal volume as well as increases in breathing frequency during severe hypoxia (8% O_2_; n = 26)^5^, including therein the Augmenting Phase (p<0.05 relative to TH-Cre for tidal volume and p<0.0001 for breathing frequency), Roll Off (p<0.001 relative to TH-Cre) and the Sustained Phase (p<0.0001 relative to TH-Cre).

Taken together these findings strongly suggest that LKB1 and AMPK facilitate the HVR and oppose respiratory depression during hypoxia. However, outcomes indicate that those catecholaminergic circuit mechanisms that mediate hypoxia-evoked increases in tidal volume are afforded greater protection from the impact of LKB1 and AMPK deficiency than those delivering increases in breathing frequency.

### *Lkb1* deletion causes marked ventilatory instability, apnoea and Cheyne-Stokes-like breathing during hypoxia

Unlike our previously reported findings in mice with *AMPKα1+α2* deletion^5^, average measures (excluding apnoeas) for *Lkb1* knockouts indicated significant augmentation rather than attenuation of increases in tidal volume during severe hypoxia, as mentioned above (Fig 6b). Closer inspection revealed that attenuation of the HVR in *Lkb1* knockouts during exposure to severe hypoxia was associated with periods of Cheyne-Stokes-like breathing (CSB), where tidal volume exhibited marked sinusoidal variations with time (Fig 7a-b**; Supplementary Movie 2**). CSB in *Lkb1* knockout mice was generally separated by periods of hypoventilation interspersed with frequent, prolonged apnoeas (≤4s). Unlike CSB, hypoventilation and apnoea were observed during mild and severe hypoxia. Increases in apnoea frequency (p<0.05 for 12% O_2_ and p<0.0001 for 8% O_2_), apnoea duration (p<0.0001) and apnoea duration index (frequency x duration; p <0.0001) were all significantly greater than for controls (TH-Cre; Fig 7c-e). As we have observed with respect to minute ventilation these measures also increased in a manner directly related to the severity of hypoxia. Moreover, CSB and increases in apneoa frequency and duration observed during severe hypoxia were completely reversed by hypercapnic hypoxia (Fig 7aIII**; Supplementary Movie 3**), likely due to improved oxygen supply consequent to increases in ventilation (see below). Therefore, the appearance of CSB likely accounts for measured increases in tidal volume for *Lkb1* knockout mice relative to controls, despite the appearance of frequent prolonged apnoeas and lengthy intervening periods of pronounced hypoventilation, that are highlighted by Poincaré plots of inter-breath interval (BBn) versus subsequent inter-breath interval (BBn+1; **Supplementary Fig 7 and 8**).

**Figure 7.**
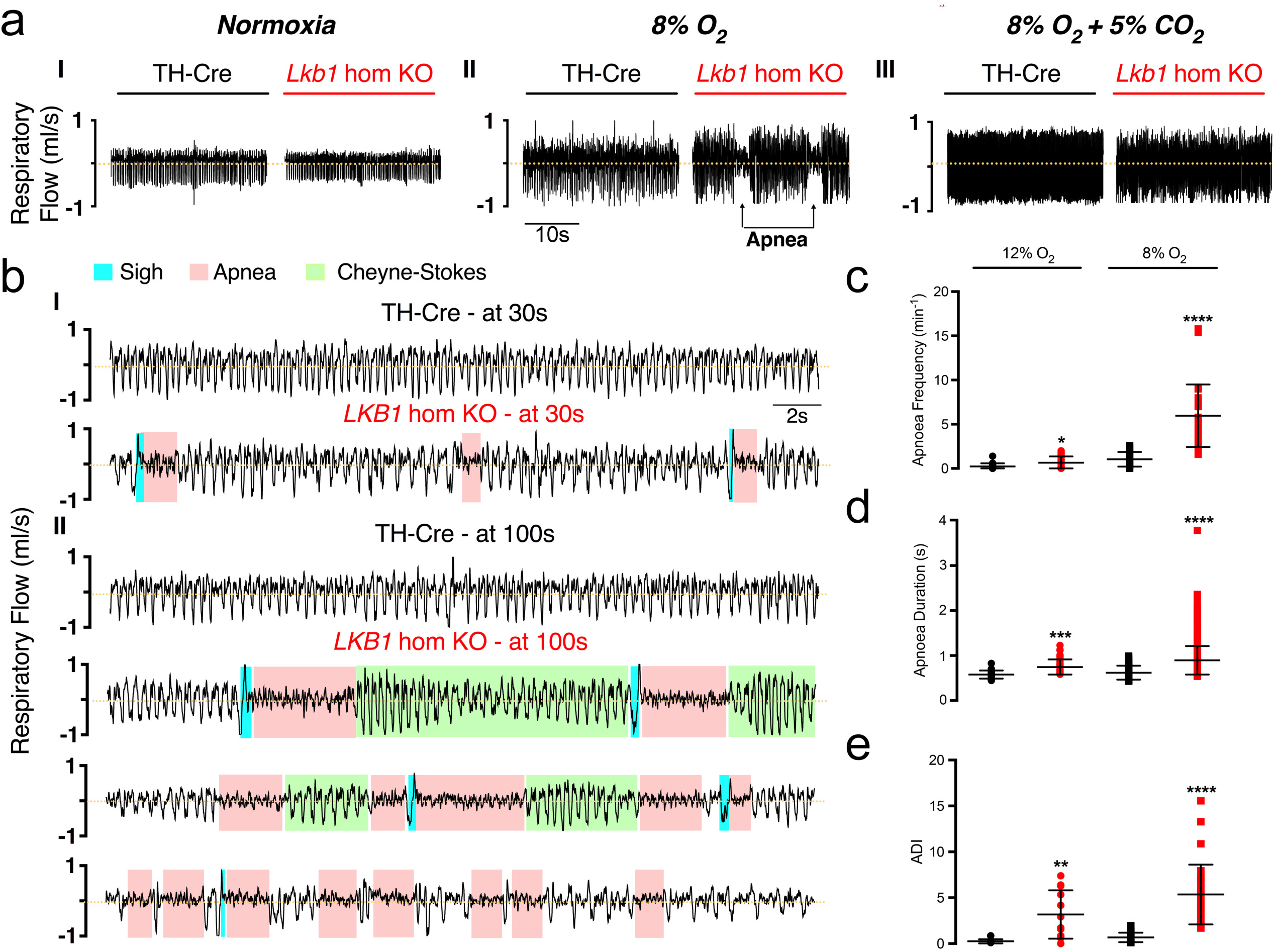
Conditional deletion of *Lkb1* in tyrosine hydroxylase expressing cells precipitates hypoventilation, apnoea and Cheyne-Stokes-like breathing during severe hypoxia. a, Example records of ventilatory activity from TH-Cre and conditional *Lkb1* homozygous knockout mice (*Lkb1* hom KO) during (I) normoxia (21% O_2_), (II) hypoxia (8% O_2_) and (III) hypoxia with hypercapnia (8% O_2_ + 5% CO_2_), that were obtained using whole body plethysmography. b(I-II), Typical ventilatory records for TH-Cre and conditional *Lkb1* hom KO mice on an expanded time scale at the indicated time points during exposures to severe hypoxia (8% O_2_). Dot plots show mean±SEM for (c) apnoeic frequency, (d) apnoea duration and (e) apnoea-duration index (frequency x duration) for TH-Cre (black; 12% O_2_, n = 19 independent experiments; 8% O_2_, n = 24 independent experiments) and conditional *Lkb1* hom KO mice (red; 12% O_2_, n = 17 independent experiments; 8% O_2_, n = 29 independent experiments) during exposures to 12% O_2_, 8% O_2_ and 8% O_2_ + 5% CO_2_. *=p<0.05, **=p<0.01, ****=p< 0.0001.

In this context, it is interesting to note that hypoxia-evoked CSB in *Lkb1* knockouts occurred irrespective of whether they were preceded by spontaneous or post-sigh apnoeas (Fig 7b). Moreover frequent and prolonged spontaneous and post-sigh apnoeas were also observed in *AMPK-α1+α2* knockouts, where CSB is absent during 5min (**Supplementary Movie 4**)^5^ or even 10min (**Supplementary Fig 9**) exposures to severe hypoxia. Therefore, if sighs were triggered by hypoxia at a given threshold ^34^, central hypoxia is likely no more severe for *Lkb1* when compared to *AMPKα1+α2* knockouts. CSB is thus most likely a consequence of LKB1 deficiency in type I cells and downstream catecholaminergic cardiorespiratory networks.

### Conditional *Lkb1* deletion slows the ventilatory response to hypercapnia and hypercapnic hypoxia

The ventilatory response to hypercapnic hypoxia (8% O_2_ + 5% CO_2_; n = 15) in *Lkb1* knockouts was attenuated, but only during the rising phase (Fig 8a; p<0.01 relative to TH-Cre, n = 17). In short, *Lkb1* deletion slowed the rising phase of the response to this stimulus but did not affect the peak achieved. It is conceivable that the slow rise in this response may result from the residual attenuation of ventilatory responses to hypoxia that are not compensated for by increased central hypercapnic ventilatory drive. However, the rise in minute ventilation during exposure to hypercapnia alone (5% CO_2_; n = 20) was also slower for *Lkb1* knockouts relative to controls (p<0.05 relative to TH-Cre, n = 20), but thereafter achieved an equivalent magnitude (Fig 8b) through increases in both respiratory frequency and tidal volume **(Supplementary Fig 10**). By contrast, mice with *AMPKα1+α2* deletion exhibited no such delay in onset of hypercapnic (n = 23) or hypoxic hypercapnic (n = 22) ventilatory responses^5^ (Fig 8a-b). The most likely explanation for these findings, therefore, is loss of carotid body chemoafferent input responses in *Lkb1* knockouts. This is in accordance with our aforementioned finding that hypoxia- and hypercapnia-evoked increases in afferent discharge were virtually abolished in carotid bodies from *Lkb1* knockouts and the generally held view that carotid body chemoafferent input responses drive the augmenting phase of the HVR ^31, 35^.

**Figure 8.**
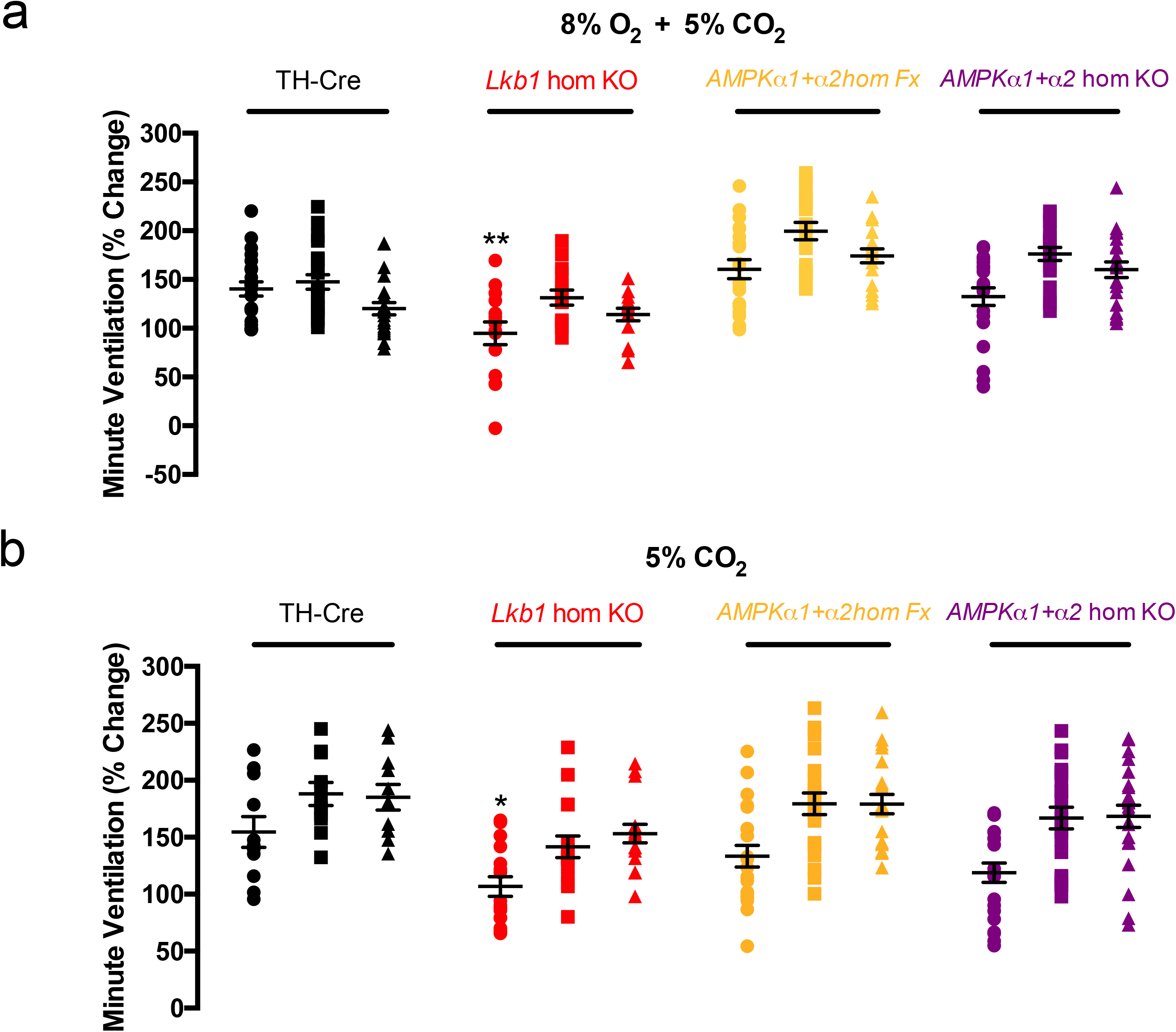
Conditional deletion of *Lkb1* in tyrosine hydroxylase expressing cells markedly slows the hypercapnic ventilatory response. Dot plots show mean±SEM for increases in minute ventilation at ∼30s, 100s and 300s during exposures to (a) hypercapnic hypoxia (5% CO_2_ + 8%O2) and (b) hypercapnia (5% CO_2_) for TH-Cre (black; 8% O2 + 5% CO_2_ n = 17 independent experiments; 5% CO2, n = 20 independent experiments), conditional *Lkb1* homozygous knockout mice (*Lkb1* hom KO, red; 8% O2 + 5% CO2, n = 15 independent experiments; 5% CO_2_, n = 20 independent experiments), *AMPKα1*+*α2* homozygous floxed mice (*AMPKα1*+*α2* hom Fx, beige; O2 + 5% CO_2_ n = 20 independent experiments; 5% CO2, n = 20 independent experiments) and *AMPKα1*+*α2* homozygous knockout mice (*AMPKα1*+*α2* hom KO, purple, O2 + 5% CO_2_ n = 22 independent experiments; 5% CO2, n = 23 independent experiments). *=p<0.05, **=p<0.01.

### The rank order of severity for attenuation of carotid body activation and attenuation of the HVR by LKB1 and AMPK is different

Peak afferent discharge from the carotid body during hypoxia remained unaffected following dual *AMPKα1+α2* deletion^5^, while hypomorphic expression of LKB1 modestly attenuated increases afferent fibre discharge from the carotid body during hypoxia and *Lkb1* deletion virtually abolished carotid body afferent discharge during normoxia and hypoxia. By contrast, the Sustained Phase of the HVR during severe hypoxia was modestly attenuated by hypomorphic expression of *Lkb1*, markedly attenuated by homozygous *Lkb1* deletion but most severely attenuated by *AMPKα1+α2* deletion. In short, the rank order by degree of inhibition of peak carotid body afferent fibre discharge during hypoxia on the one hand and the sustained phase of the HVR on the other is different (**Supplementary Fig 11**).

## Discussion

The present study identifies an essential role for LKB1 in establishing carotid body function and chemosensitivity, where the level of LKB1 expression determines a set-point about which carotid body afferent discharge is modulated by hypoxia and hypercapnia. This strongly suggests that the metabolic signalling pathway(s) that mediates the response of carotid body type I cells to hypoxia cannot be attenuated without affecting CO_2_ sensitivity. Moreover, we have uncovered a divergence in dependency on LKB1 and AMPK between the carotid body on the one the hand and the hypoxic ventilatory response (HVR) on the other (Fig 9).

**Figure 9.**
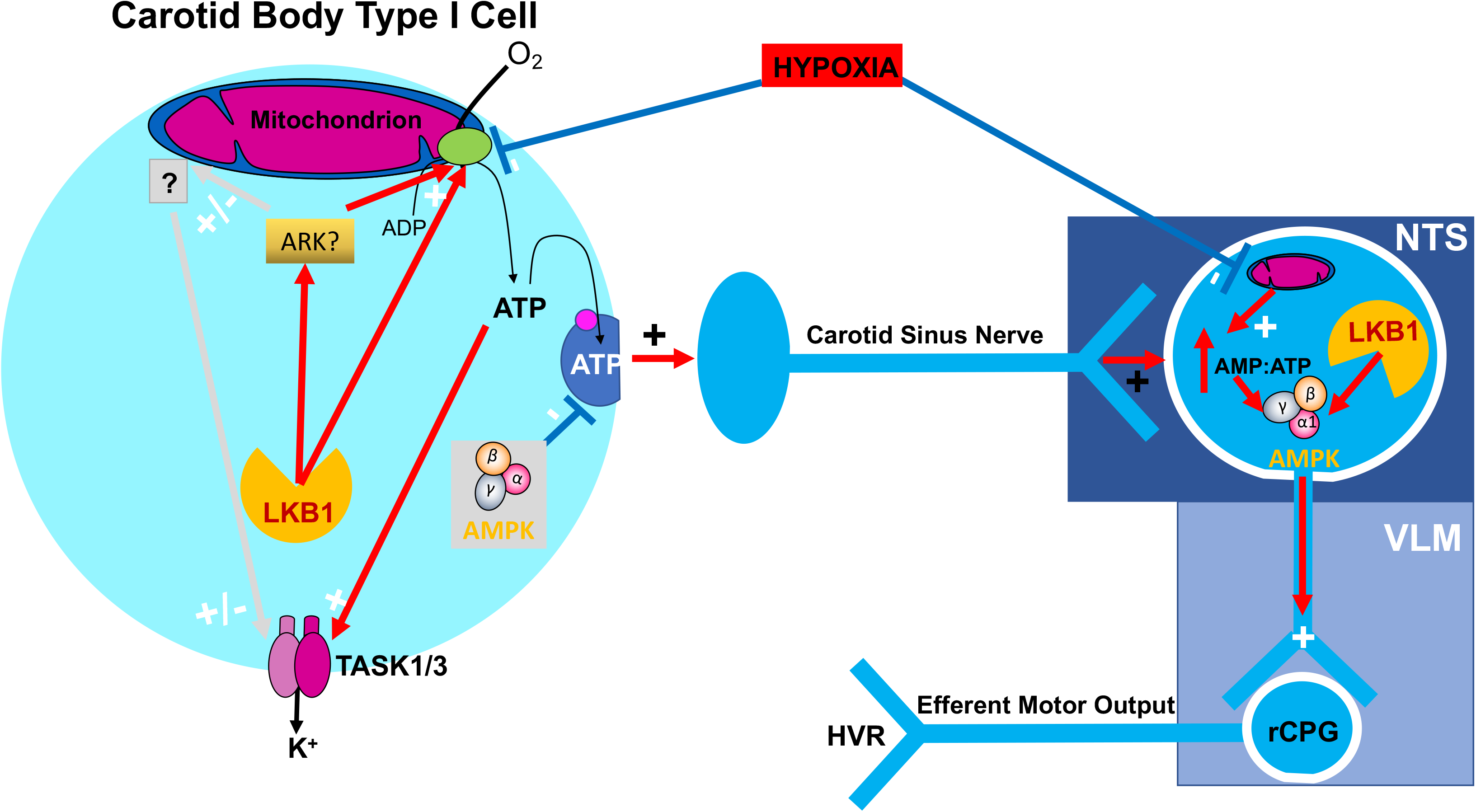
Graphical abstract showing the divergent pathways by which LKB1 and AMPK may coordinate the hypoxic ventilatory response: LKB1, liver kinase B1; AMPK, AMP-activated protein kinase; ARK, AMPK-related kinase; NTS nucleus tractus solitarius; VLM, ventrolateral medulla.

Deletion of LKB1, but not AMPK, in type I cells attenuated basal carotid body afferent discharge during normoxia and virtually abolished carotid body afferent input responses during hypoxia and hypercapnia, the latter of which was not thought to be determined by changes in mitochondrial metabolism ^26–29^. Carotid bodies of mice that are ∼90% hypomorphic for LKB1 expression in all cells (homozygous *Lkb1* floxed ^23^) also exhibited significantly attenuated (∼50%) peak carotid body afferent discharge during hypoxia. Accordingly, homozygous *Lkb1* deletion abolished hypoxia-evoked cytoplasmic calcium transients in type I cells. Paradoxically, however, in type I cells from mice that were ∼90% hypomorphic for LKB1 expression, hypoxia-evoked calcium transients were not only retained but augmented relative to controls. This suggests that the level of LKB1 expression determines a set point about which type I cells are activated by hypoxia and hypercapnia, and that membrane depolarisation and exocytotic transmitter release are differentially sensitive to this.

It has been proposed that afferent discharge during hypoxia could, at least in part, be initiated by falls in type I cell cytoplasmic ATP ^36^ (although other mechanisms have also been considered ^37^), which could trigger membrane depolarisation^36^ and consequent exocytotic release of vesicular ATP to induce increases in afferent discharge ^38, 39^. It is intriguing to note, therefore, that previous studies on the effects of *Lkb1* deletion have identified changes mitochondrial activities and consequent reductions in ATP levels and increases in AMP:ATP and ADP:ATP ratios in a variety of cell types, including skeletal muscle ^23^, cardiac muscle ^24, 40^, pancreatic beta cells ^41 42^, regulatory T Cells^43^ and MIN6 cells ^41^. More importantly still, it has been demonstrated that ATP levels are lower in cardiac muscle from hypomorphic *Lkb1* floxed mice under normoxia, lower still in hearts from mice with cardiac-specific *Lkb1* deletion and that in each case ATP levels decline further during ischaemia ^24^. Thus, LKB1 may maintain in an expression-dependent manner the capacity for ATP synthesis within most cells, including therein carotid body type I cells where ATP deficiency might ultimately impact on the capacity for uptake of ATP by synaptic vesicles and/or exocytotic release of ATP. This view gains indirect support from our finding that *Lkb1* deletion, but not hypomorphic expression of LKB1, showed signs of reducing basal afferent discharge during normoxia, while peak afferent discharge was mildly attenuated by hypomorphic expression of LKB1 (∼50%) and virtually abolished by *Lkb1* deletion. Further indirect support is provided by a previous study on rat carotid bodies, which showed that exocytotic release of adenosine represents the principal transmitter contributing to carotid body afferent discharge during mild hypoxia, while ATP acts as the principal transmitter driving afferent discharge during severe hypoxia (i.e., peak discharge frequency) ^44^. Although counter-intuitive, a lowering of ATP levels could also explain why hypoxia-evoked calcium transients were augmented in carotid body type I cells with hypomorphic expression of LKB1 yet blocked by homozygous *Lkb1* deletion, if mid-range reductions in basal ATP levels confer increased sensitivity of TASK1/3 channels to inhibition by hypoxia as previously proposed by others^36^, without greatly compromising the capacity for exocytotic ATP release. Further reductions in ATP availability upon homozygous *Lkb1* deletion could then ultimately render these ATP-sensitive TASK1/3 channels inactive, abolish hypoxia-response coupling in type I cells ^16, 36, 45^ and greatly reduce vesicular uptake and exocytotic ATP release^38, 39^. This would also impact on type I cell activation by hypercapnia, due to the fact that this too is in great part mediated by inhibition of TASK1/3 channels through acidosis and consequent induction of exocytotic release of ATP, albeit in a manner independent of mitochondrial oxidative phosphorylation ^30^. Indirect support for this proposal may be taken from the previous finding of others that inhibition of mitochondrial oxidative phosphorylation (oligomycin and antimycin A) in cat carotid bodies in-situ blocked responses to hypoxia and enhanced responses to hypercapnia *in-vivo*^30^. If LKB1 deficiency did not reduce ATP availability, then one would expect similarly augmented changes in afferent discharge in response to hypercapnia rather than the decreases reported here. It is also notable that *Lkb1* deletion in pancreatic beta cells is associated with lower glucose-induced ATP accumulation, enhanced membrane excitability and increased glucose-stimulated insulin release ^41, 42^. That said it is possible that LKB1 deficiency may increase or reduce other metabolic intermediates that might also impact type I cell responses to hypercapnia and thus O_2_:CO_2_ stimulus interaction ^27, 46^. Either way, the precise AMPK-independent mechanism(s) by which LKB1 may “rewire cell metabolism” remains to be determined. However, it is intriguing to note that LKB1 can coordinate glucose homeostasis ^41, 47, 48^ and mitochondrial function through either a direct action^41, 49, 50^, or indirectly through constitutive phosphorylation of one or more of the eleven AMPK-related kinases it regulates^13, 51, 52^.

In line with the above, the HVR was attenuated in hypomorphic *Lkb1* floxed and homozygous *Lkb1* knockout mice in a manner related to the degree of LKB1 deficiency. By contrast, the HVR remained unaffected in global *Camkk2* knockouts. In short, LKB1 and AMPK signalling pathways are critical to the maintenance of breathing and oxygen supply during hypoxia^53^, and act in concert to oppose ventilatory depression, hypoventilation and apnoea^5^.

We previously reported that dual *AMPKα1+α2* deletion in catecholaminergic cells blocked the HVR, with no discernable effect on carotid body afferent input responses to hypoxia^5^. The data presented here confirm this finding and show that carotid body CO_2_ sensitivity remained unaltered following *AMPKα1+α2* deletion. Therefore, our present findings support the idea that severe hypoventilation and apnoea observed during hypoxia in *AMPKα1+α2* knockout mice is due to dysfunction of central respiratory networks rather than any depletion of carotid body activity.

It is evident that *Lkb1* deletion in catecholaminergic neurons attenuated all phases of the HVR during mild (12% O_2_) and severe (8% O_2_) hypoxia, to a greater degree than with hypomorphic expression of LKB1 but less so than previously observed during severe hypoxia following *AMPKα1+α2* deletion, even though carotid body afferent input responses were retained in *AMPKα1+α2* knockouts^5^. In this respect, it is interesting to note that deficits in minute ventilation were evident for *Lkb1* floxed mice during the late Sustained Phase but not the Augmenting Phase of the HVR during severe hypoxia. This is consistent with the effect of *AMPKα1+α2* deletion during mild hypoxia. Outcomes for *Lkb1* floxed mice and *AMPKα1+α2* knockouts therefore add further weight to our proposal that they exert an inhibitory effect downstream of chemoafferent input responses, if one accepts the view that increases in carotid body afferent discharge drive the Augmenting Phase of the HVR ^31, 35^ while direct modulation by hypoxia of brainstem respiratory networks aids maintenance of the HVR in the longer term ^1, 5, 32, 35, 54, 55^. Further support for this proposal is provided by the rank order by severity of hypoxic ventilatory and carotid body dysfunction, respectively. During severe hypoxia the HVR is inversely related to the degree of LKB1 deficiency but was most markedly inhibited following *AMPKα1+α2* deletion^5^, despite the fact that carotid body afferent discharge during hypoxia (and hypercapnia) was attenuated in a manner directly related to the degree of *Lkb1* deletion but remained unaffected following *AMPKα1+α2* deletion.

When taken together these data strongly suggest that LKB1 determines, independent of AMPK, a set-point about which carotid body afferent input responses are delivered during hypoxia and provide further strong support for our previous proposal that LKB1-AMPK signalling pathways facilitate the HVR at the brainstem. In this way LKB1 and AMPK may exert independent influences on peripheral ^27, 46^ and central^31, 35, 55, 56^ stimulus interactions. Accordingly, the rising phase of the hypercapnic and hypoxic-hypercapnic ventilatory responses was slowed in *Lkb1* knockouts, while by contrast the peak of the Sustained Phase of both responses remained unaltered despite marked attenuation of afferent input responses to hypercapnia.

This is a major point because homozygous *Lkb1* deletion led to marked reductions in breathing frequency (excluding apnoeas) that were coupled with erratic “augmentation” of tidal volume responses during severe hypoxia, consequent to induction of periodic Cheyne-Stokes-like breathing patterns. Cheyne-Stokes-like breathing patterns were never observed in mice with either ∼90% global hypomorphic LKB1 expression or *AMPKα1+α2* deletion in catecholaminergic cells. A critical distinguishing factor in this respect may therefore be the block by *Lkb1* deletion of not only carotid body afferent input responses to hypoxia and hypercapnia, but also concomitant attenuation of downstream hypoxia-responsive circuit mechanisms. This suggests that Cheyne-Stokes breathing may occur consequent to energy crises in peripheral and central catecholaminergic respiratory control networks. That said, others have proposed that Cheyne-Stokes breathing is triggered by hyperactivity of carotid bodies and thus augmented afferent input responses when associated with heart failure ^57^. One possible explanation for these contradictory observations could be that enhanced carotid body afferent input responses after heart failure occur consequent to central metabolic crisis that results in abject failure of both central integration of afferent inputs and efferent ventilatory output. In other words, increases in controller gain within the central respiratory networks could trigger Cheyne-Stokes breathing by enhancing the sensitivity to, and thus the degree of activation of central CO_2_-sensing neurons during hypercapnia ^31^ consequent to hypoventilation and apnoea during hypoxia. Consistent with this view, others have proposed that Cheyne-Stokes breathing may be caused by enhanced hypercapnic ventilatory responses driven by instability within respiratory networks consequent to augmented chemoreflex gain, prolonged feedback delay ^58^ and/or enhanced central controller gain ^59^.

The more extreme patterns of non-rhythmic (ataxic) ventilation observed for *AMPKα1+α2* knockouts ^5^ may thus be avoided. While unlikely it is also conceivable that retention by *Lkb1* knockouts of greater capacity for rhythmic ventilation during hypoxia could be conferred by residual allosteric AMPK activation by AMP in central hypoxia-responsive respiratory networks^60^, where falls in cellular ATP supply would be associated with ADP accumulation and consequent increases in the AMP:ATP ratio via the adenylate kinase reaction. This could conceivably maintain oscillating central respiratory drive in a manner triggered periodically once a given severity of central hypoxia is breached. That said, central hypoxia is likely no more severe for *Lkb1* when compared to *AMPKα1+α2* knockouts because: (1) hypoxia-evoked sighs^34, 61^ were observed in *Lkb1* and *AMPKα1+α2* knockouts; (2) apnoeas were shorter and less frequent for *Lkb1* knockouts when compared to *AMPKα1+α2* knockouts ^5^; (3) Cheyne-Stokes-like breathing between apnoeas would periodically raise oxygen supply in *Lkb1* knockouts.

In conclusion, the present study reveals that the level of LKB1 expression is essential for establishing carotid body function and for initiating the HVR. In this respect LKB1 and AMPK provide for hierarchical control of the hypoxia-responsive respiratory network (Fig 9). Firstly, the level of LKB1 expression determines, independent of AMPK, a set-point about which carotid body afferent input responses are evoked during hypoxia and hypercapnia, rather than contributing to oxygen-sensing *per se*. Thereafter LKB1-AMPK signalling pathways likely govern coincidence detection and signal integration within an hypoxia-responsive circuit downstream of the carotid bodies, that encompasses, at the very least, the brainstem nucleus of the solitary tract ^5^. Afferent input responses and brainstem hypoxia could thereby determine, each in part, the set-point about which AMPK and thus brainstem respiratory networks are activated during hypoxia. Subsequently, AMPK-dependent modulation of cellular metabolism ^60^, ion channels ^62, 63^ and thereby neuronal activities ^64, 65^ may facilitate afferent inputs and thus efferent outputs leading to increases in ventilatory drive during hypoxia. Consequently, LKB1 and/or AMPK deficiency may contribute to central sleep apnoea associated with metabolic syndrome-related disorders ^66^, ascent to altitude ^67^ and apnoea of prematurity ^68^. By contrast, Cheyne-Stokes breathing and central sleep apnoea ^58^ associated with heart failure ^57^ may be conferred by LKB1 deficiency and/or metabolic crises across peripheral and central hypoxia-responsive respiratory networks. Further studies are therefore warranted to elucidate the downstream AMPK-independent targets by which LKB1 establishes carotid body function.

## Methods

Experiments were approved by local ethical review committees and the University Director of Veterinary Services at the University of Edinburgh, and by the UK Home Office (Science). All procedures were covered by a UK Home Office Project Licence (PBA4DCF9D). All genetically modified mice tested here were bred on a C57 Black 6 (C57/BL6) background. Furthermore, all studies complied with the regulations of the United Kingdom Animals (Scientific Procedures) Act of 1986.

### Breeding of mice, genotyping and single cell PCR

Standard approaches were used for breeding of mice and brother/sister mating was avoided. All mice studied were between 3-12 months of age.

Because global deletion of the gene encoding LKB1 (*Stk11, Lkb1*) or dual deletion of the genes encoding AMPKα1 (*Prkaa1*) and AMPKα2 (*Prkaa2*) is embryonic lethal, we employed knockdown and/or conditional deletion strategies. For *Lkb1* deletion we used floxed mice in which exons 5-7 of this gene had been replaced by a cDNA cassette encoding equivalent exon sequences where exon 4 and the cDNA cassette were flanked by loxP sequences, which in their own right deliver ∼90% global knockdown of LKB1 expression ^23^. For *AMPKα1+α2* deletion critical exons of the *AMPKα1* and *AMPKα2* genes were flanked by loxP sequences ^69^. Each floxed mouse line was crossed, as previously described ^5^, with mice expressing Cre recombinase under the control of the tyrosine hydroxylase (TH) promoter (Th-IRES-Cre; EM:00254), providing for gene deletion in all catecholaminergic cells inclusive of those cells that constitute the hypoxia-responsive respiratory network from carotid body ^70^ to brainstem ^71^. Transient developmental expression of TH does occur in disparate cell types that do not express TH in the adult ^72^, such as dorsal root ganglion cells and pancreatic islets, but these do not contribute to the acute HVR. We previously confirmed restriction of Cre expression to TH-positive cells in the adult mouse by viral transfection of a Cre-inducible vector carrying a reporter gene ^5^. Therefore, our approach overcomes embryonic lethality and allows, unforeseen ectopic Cre expression aside, for greater discrimination of circuit mechanisms than would be provided for by global knockouts. The role of CaMKK2 in the HVR was determined by assessing mice with global deletion of the corresponding gene (*Camkk2*) ^22^.

Male Lkb1^flx/flx^ mice are infertile. To overcome this issue female Lkb1^flx/flx^ mice were crossed with heterozygous male TH-Cre^+/-^ mice. Heterozygous males of the Lkb1^flx/wt^ Cre^+/-^ genotype were then backcrossed with female homozygous Lkb1^flx/flx^ Cre^+/-^ mice to obtain the required Lkb1^flx/flx^ Cre^+/-^ mice to study. Wild type or floxed *Lkb1* alleles were detected using two primers, p200, 5’-CCAGCCTTCTGACTCTCAGG-3’ and p201, 5’-GTAGGTATTCCAGGCCGTCA-3’. For the detection of Cre recombinase we employed: TH3, 5’-CTTTCCTTCCTTTATTGAGAT-3’, TH5, 5’-CACCCTGACCCAAGCACT-3’ and Cre-UD, 5’-GATACCTGGCCTGGTCTCG-3’. As homozygous *Lkb1* floxed mice are hypomorphic, exhibiting ∼90% lower LKB1 expression than *Lkb1* wild type littermates ^23^, we used as controls mice that express Cre via the tyrosine hydroxylase promoter (TH-Cre).

For deletion of the gene that encodes CaMKK2 (*CamKK2*) wild type alleles were detected using two primers, KKBeta1, 5’CAGCACTCAGCTCCAATCAA3’, and KKBeta2, 5’GCCACCTATTGCC TTGTTTG3’.

Lastly, we used two primers for each AMPK catalytic subunit: α1-forward: 5’ TATTGCTGCCATTAGGCTAC 3’, α1-reverse: 5’ GACCTGACAGAATAGGATATGCCCAACCTC 3’; α2-forward 5’ GCTTAGCACGTTACCCTGGATGG 3’, α2-reverse: 5’ GTTATCAGCCCAACTAATTACAC 3’.

We detected the presence of wild-type or floxed alleles *and* Cre-recombinase by PCR. The PCR protocol used for all genotype primers was: 92°C for 5min, 92°C for 45s, 56°C for 45s, 72°C for 60s, and 72°C for 7min for 35 cycles and then 4°C as the holding temperature. 15µl samples were run on 2% agarose gels with 10µl SYBR®Safe DNA Gel Stain (Invitrogen) in TBE buffer against a 100 bp DNA ladder (GeneRuler^TM^, Fermentas) using a Model 200/2.0 Power Supply (Bio-Rad). Gels were imaged using a Genius Bio Imaging System and GeneSnap software (Syngene).

### Type I cell isolation

Carotid bodies were incubated at 37°C for 25-30min in isolation medium consisting of: 0.125mg/ml Trypsin (Sigma), 2.5mg/ml collagenase Type 1 (Worthington) made up in low Ca^2+^/low Mg^2+^ HBSS. During this incubation the carotid bodies were separated from the associated patch of artery. The carotid bodies were then transferred to low Ca^2+^/low Mg^2+^ HBSS containing trypsin inhibitor (0.5mg/ml) for 5min at room temperature, and then to 2ml of pre-equilibrated (95% air, 5% CO_2_, 37°C) growth medium (F-12 Ham nutrient mix, 10% fetal bovine serum, 1% penicillin/streptomycin). The medium containing the carotid bodies was centrifuged and the pellet re-suspended in 100µl of growth medium. Carotid bodies were then disrupted by triturating using fire polished Pasteur pipettes, and type I cells used within 4 hr.

### Confocal and Immunofluorescence imaging

To aid confirmation that *Lkb1* deletion had been induced in carotid body type I cells, mice with TH-Cre driven gene deletion were crossed with mice engineered for Cre-dependent expression of tdTomato (excitation 555 nm, emission 582 nm) from the Rosa26 locus. These mice were deeply anaesthetised using 2g/kg Pentobarbital Sodium (Merial) and carotid bifurcations containing the carotid body tissue dissected out. Bifurcations were briefly washed in ice-cold saline, fixed in 4% paraformaldehyde in 0.1M phosphate buffer (PB; pH 7.4), post-fixed, and stored in 30% sucrose in 0.1M PB at 4°C. 5µm sections of the bifurcations were cut using a cryostat, collected on glass slides and air dried before being rinsed in 0.1M phosphate buffered saline (PBS) and glass coverslipped. Confocal z sections were acquired using a Nikon A1R + confocal system via a Nikon Eclipse Ti inverted microscope with a Nikon Apo 63Å∼ λS DIC N2, 1.25 n.a. oil immersion objective (Nikon Instruments Europe BV, Netherlands). Image processing was carried out using Imaris (Bitplane, Oxford Instruments, UK) and Image J (Rasband WS. ImageJ, U.S. National Institutes of Health, Bethesda, MD, USA, imagej.nih.gov/ij/, 1997–2012).

Additionally, carotid body type I cells were isolated and processed for immunocytochemistry. Briefly, slides were washed in 0.1M phosphate buffered saline, incubated overnight in anti-TH (mouse; 1:1000 dilution; Merck Millipore MAB318) primary antibodies diluted in 2% v/v normal serum in 0.1M PB-T (0.3% v/v Triton^TM^ X-100; Sigma), rinsed 3x for 5min in 0.1M PBS and incubated in fluorescent secondary antibodies (Alexa Fluor^®^ 488; goat anti-mouse; 1:750 dilution; Thermo Fisher A-11034) for 2hr at room temperature. Slides were washed again, followed by incubation with DAPI (1 μg/ml) for 5min at room temperature, 3x 5min washed with 0.1M PBS and glass coverslipped.

### Single-cell end-point PCR

For single cell amplification, GoTaq DNA Polymerase (Promega) was added to 2-5μl of cDNA obtained from each single carotid body type I cell from wildtype and transgenic mice as well as the wildtype adrenomedullary chromaffin cells. To ensure the validity that the collected cells were indeed carotid body type I cells and adrenomedullary chromaffin cells, primers obtained from Qiagen were used to detect the expression of tyrosine hydroxylase (QuantiTect Primer Assay, QT00101962) with an expected band length of 92bp. The only cells considered for the expression studies were those that positively expressed tyrosine hydroxylase and where the negative controls were clean. Primers designed by Qiagen for LKB1 were not used as they detect area of the genes that are not within the floxed loxP sites, which may result in false positives appearing if mRNA transcript is still produced regardless of whether the targeted domain has been excised. Accordingly, primers were designed by using Primer-BLAST (NCBI) to detect a region that is known to be excised: FWD: 5’GCTCATGGGTA CTTCCGCCAGC 3’; REV:5’AGCAGGTTGCC CGGCTTGATG 3’. 15μl samples along with a 100bp DNA Ladder (GeneRulerTM, Fermentas) were run on a 2% agarose gel made with SYBR®Safe DNA gel stain (Invitrogen). Gels were then imaged using a Genius Bio Imaging System and GeneSnap Software (Syngene).

### Quantitative RT-PCR

RNA was extracted, quantified and reverse transcribed as described above. For qPCR analysis, 2.5µl of cDNA in RNase free water was made up to 25µl with FastStart Universal SYBR Green Master (ROX, 12.5µl, Roche), Ultra Pure Water (8µl, SIGMA) and forward and reverse primers for LKB1. The sample was then centrifuged and 25µl added to a MicroAmp^TM^ Fast Optical 96-Well Reaction Plate (Greiner bio-one), the reaction plate sealed with an optical adhesive cover (Applied Biosystems) and the plate centrifuged. The reaction was then run on a sequence detection system (Applied Biosystems) using AmpliTaq Fast DNA Polymerase, with a 2min initial step at 50°C, followed by a 10min step at 95°C, then a 15s step at 95°C which was repeated 40 times. Then a dissociation stage with a 15s step at 95°C followed by a 20s at 60°C and a 15s step at 95°C. Negative controls included control cell aspirants for which no reverse transcriptase was added, and aspiration of extracellular medium and PCR controls. None of the controls produced any detectable amplicon, ruling out genomic or other contamination.

### Calcium imaging

Type I cells were incubated in the standard perfusate with 4µM Fura-2 AM (Molecular Probes) at room temperature, washed and then placed in a temperature regulated perfusion chamber on a Nikon Diaphot 300 inverted phase contrast microscope. Cells were perfused with a solution consisting of (mM): NaCl (117), KCl (4.5), MgCl_2_ (1), CaCl_2_ (2.5), NaHCO_3_ (23), Glucose (10), pH adjusted to 7.4 using 5% CO_2_. For normoxia perfusate was bubbled with 5% CO_2_ 95% air (∼150mmHg). For hypoxia perfusate was bubbled with 5% CO_2,_ 95 % nitrogen (∼20mmHg).

### Extracellular recordings of carotid sinus nerve activity

Single fibre chemoafferent activity was amplified, filtered and recorded using a 1401 interface running Spike 2 software (Cambridge Electronic Design). Single- or few-fibre chemoafferent recordings were made from carotid bifurcations held in a small volume tissue bath (36-37°C), and superfused with gassed (95% O_2_ and 5% CO_2_), bicarbonate-buffered saline solution (composition (mM): 125 NaCl, 3 KCl, 1.25 NaH_2_PO_4_, 5 Na_2_SO_4_, 1.3 MgSO_4_, 24 NaHCO_3_, 2.4 CaCl_2_, pH 7.4). A standard O_2_ electrode (ISO2; World Precision Instruments) was placed in the superfusate system at the point of entry to the recording chamber in order to continuously record the superfusate *P*O_2_. Flow meters with high precision valves (Cole Palmer Instruments) were used to equilibrate the superfusate with a desired gas mixture. Basal single fibre activity was monitored at a superfusate *P*O_2_ of 200mmHg and a PCO_2_ of 40mmHg. This *P*O_2_ is slightly lower than that previously used for the rat carotid body ^73^ to take in account the smaller size of this organ in the mouse (and thus a smaller diffusion distance). At this superfusate *P*O_2_, the basal frequency in TH-Cre single/few fibres was consistent with that reported *in vivo* in other rodents^74^ and so we interpret this *P*O_2_ to have not been excessively hyperoxic.

To induce responses to hypoxia, the superfusate *P*O_2_ was slowly reduced to a minimum of 40mmHg or was reversed prior to this when the chemoafferent response had stabilised or had begun to diminish. The single fibre chemoafferent discharge frequency was plotted against the superfusate *P*O_2_ over a desired range of superfusate *P*O_2_ values. To produce the hypoxic response curves, the data points were fitted to an exponential decay curve with offset, as shown below:

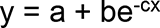

For the above equation, y is the single fibre discharge frequency in Hz, x is the superfusate *P*O_2_ in mmHg, a is the discharge frequency as the *P*O_2_ tends to infinity (offset), b is the discharge frequency when the *P*O_2_ is 0mmHg (minus the offset) and c is the exponential rate constant. Comparison of the exponential rate constants allowed for determination of any alteration in the rate of increase in chemoafferent frequency per mmHg reduction in the superfusate *P*O_2_, upon hypoxic response initiation. Furthermore, for any given discharge frequency, the corresponding *P*O_2_ could be calculated using the inverse function of the exponential decay curve, as shown below:

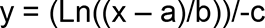

y is the *P*O_2_ in mmHg, x is the single fibre discharge frequency in Hz and a,b and c are constants as above. Specifically, superfusate *P*O_2_ levels were compared when the single fibre chemoafferent discharge frequency was at 5 Hz. This was chosen as it lies on the exponential region of the hypoxic response curve but is not of a magnitude at which the discharge is likely to have begun to diminish. This method was used to define any *P*O_2_ shift in the hypoxic response curve thereby providing information of a potential change in the *P*O_2_ threshold required for hypoxic response initiation. Plots of firing frequency versus superfusate *P*O_2_ were fitted by non-linear regression (GraphPad Prism 6).

Chemoafferent responses to hypercapnia were induced by raising the superfusate PCO_2_ from approximately 40 mmHg (pH 7.4) to 80 mmHg (pH 7.15) at a constant PO_2_ (200 mmHg), as has been previously reported for the intact *in vitro* CB preparation^75^.

### Plethysmography

For plethysmography mice were 6-12 months of age. Both males and females were studied. We used unrestrained whole-body plethysmography, incorporating a Halcyon^TM^ low noise pneumatochograph coupled to FinePointe acquisition and analysis software with a sampling frequency of 1kHz (Buxco Research Systems, UK). All quoted values for the HVR were derived from apnoea-free periods of ventilation. All measures reported are averages of n repeats from multiple mice (C57/BL6, 5 mice; TH-Cre, 5 mice; *Lkb1* floxed, 4 mice; *Lkb1* knockout, 4 mice; CaMKK2 knockouts, 7 mice; AMPK-α1/α2 floxed, 5 mice; AMPK-α1/α2 floxed, 5 mice). Any unreliable and erratic respiratory waveforms recorded during gross un-ventilatory related body movements, i.e., sniffing and grooming, were avoided for measurements. Additionally, a rejection algorithm that was built into the plethysmography system (Buxco Electronics Inc.) identified periods of motion-induced-artefacts for omission. The patented Halycon^TM^ low noise pneumotachograph (Buxco Electronics Inc.) reduces disturbances caused by air currents from outside the chambers (i.e., fans, closing doors, air conditioners, etc.), which can disrupt or overwhelm the ventilatory airflows within the chamber.

Mice were trained by repeated bi-weekly placement in the plethysmography chamber under normoxia and without experimental interventions, so that they became accustomed to the environment. During experimental work mice were placed in a freshly cleaned plethysmography chamber (to remove scent of previous mice) for a 10-20min acclimation period under normoxia (room air) to establish a period of quiet and reliable breathing for baseline-ventilation levels (this is also indicated by a measured rejection index of 0 by the FinePointe Acquisition and Analysis Software). Mice were then exposed to hypoxia (12% or 8% O_2_, with 0.05% CO_2_, balanced with N_2_), hypoxia+hypercapnia (8% O_2_, 5% CO_2_, balanced with N_2_) or hypercapnia (21% O_2_, 5% CO_2_, balanced with N_2_) for 5min or 10min. Medical grade gas mixtures were chosen by switching a gas tap. The time for evacuation of the dead space and complete exchange of gas within the plethysmography chamber was 30s. The duration of exposure to hypoxia quoted was the actual duration of hypoxia. Apnoea was defined as cessations of breathing greater than the average duration, including interval, of 2 successive breaths (600ms) during normoxia, with a detection threshold of 0.25mmHg (SD of noise). Breathing variability was assessed by Poincaré plots and by calculating the SD of inter-breath (BB) intervals. The breathing frequency, tidal volume, and minute ventilation as derived by the FinePointe Software were also analysed for control and knockout mice. These parameters were measured as mean values taken over a 2s breathing period and not on a breath-to-breath basis. The changes in breathing frequency, tidal volume, and minute ventilation during hypoxia and/or hypercapnia were analysed as the percentage change from normoxia respective to each individual mouse. The peak of the augmenting phase was calculated from the peak value between 20-40s of the hypoxic and/or hypercapnic exposure that coincides with the peak of the rising phase. The roll off period was calculated as the lowest value between 60-140s of exposure and the sustained phase was calculated from the last 20s in the plateaued phase. A large time range was required for selection of these points as experiments were performed on unrestrained and awake animals and periods of no movement, sniffing, or grooming, were only considered.

Apnoeas were excluded from all stated measures (mean±SEM) of breathing frequency, tidal volume and minute ventilation, i.e., all quoted values were derived from apnoea-free periods of ventilation.

### Statistics and Reproducibility

Statistical comparison was completed using GraphPad Prism 6 as follows: Calcium imaging data were assessed by Student’s t test; Afferent discharge was assessed by single or 2 factor ANOVA with Bonferroni Dunn post hoc analysis and by Student’s t test; Plethysmography was assessed by one-way ANOVA with Bonferroni multiple comparison’s test and by Student’s t test; p<0.05 was considered significant. For afferent discharge and plethysmography all quoted values are for ANOVA unless stated otherwise. All data are presented as mean±SEM. All responses studied were robust to inter-animal variability and highly reproducible. Replicates for calcium imaging on isolated type I cells refer to independent studies on 8-11 different cells from at least three (3) different mice. For afferent fibre discharge replicates refer to studies on 4-10 different carotid bodies each from a different mouse. For plethysmography all measures reported are averages of n separate experiments on 4-7 mice spread over a six-month period from six months of age (C57/BL6, 5 mice; TH-Cre, 5 mice; *Lkb1* floxed, 4 mice; *Lkb1* knockout, 4 mice; CaMKK2 knockouts, 7 mice; AMPK-α1/α2 floxed, 5 mice; AMPK-α1/α2 floxed, 5 mice). To ensure as few mice as possible were used to determine differences by significance test, experiments were conducted and acquired data statistically assessed in stages by the variable criteria sequential stopping rule (SSR). In this way animal use was minimised, power maximised and the probability of type I errors kept constant.

## Data availability statement

All data generated or analysed during this study are included in this published article (and its supplementary information files).

## Acknowledgements

This work was primarily funded by a Wellcome Trust Programme Grant held by AME (WT081195MA), which also supported by subcontract the work of A.P.H and P.K. at the University of Birmingham. S.M. was supported by a University of Edinburgh PhD studentship and then by a BHF Programme Grant held by AME (RG/12/14/29885).

## Author contributions

A.M.E. conceived this study and developed the study plan in discussion with D.G.H.. A.M.E. and S.M. wrote the manuscript and made Figures 1-9. A.M.E. and S.M. developed the conditional knockout mice. S.M. and A.D.M designed and validated primers and performed genotyping. M.J.S., A.D.M., S.M. and A.M.E. performed single cell PCR. A.M.D., S.M., and A.M.E. performed plethysmography. S.M. and A.M.E. performed final analysis of respiratory data. S.M. and A.M.E. developed ROSA mice and performed confocal microscopy and immunocytochemistry. A.P.H. and P.K. performed afferent discharge and carried out data analysis blind, before final data compilation and interpretation by A.M.E. in discussion with S.M., A.P.H. and P.K.. M.L.D, A.M.D. and C.P. carried out and performed initial acquisition and analysis of calcium imaging blind, which was later compiled, tested and interpreted by A.M.E. in discussion with S.M.. All authors discussed their results and provided feedback on one or more drafts of this manuscript.

