## Supplementary Information for "LKB1 is the gatekeeper of carotid body chemo-sensing and the hypoxic ventilatory response"

**RUNNING HEAD**: LKB1 carotid bodies and the HVR

Sandy MacMillan[1+], Andrew P. Holmes [2+], Mark L. Dallas[3], Amira D. Mahmoud[1], Michael J. Shipston[1], *the late* Chris Peers, D. Grahame Hardie[4], Prem Kumar[2], A. Mark Evans[1*]

**Supplementary Movie 1 - Hypoxic ventilatory response of control (TH-Cre) mouse.**

**Supplementary Movie 2 - Hypoxic ventilatory response of homozygous *Lkb1* knockout mouse.**

**Supplementary Movie 3 - Hypercapnic hypoxic ventilatory response of homozygous *Lkb1* knockout mouse.**

**Supplementary Movie 4 - Hypoxic ventilatory response of homozygous *AMPK*$\alpha$*1*+$\alpha$*2* knockout mouse.**

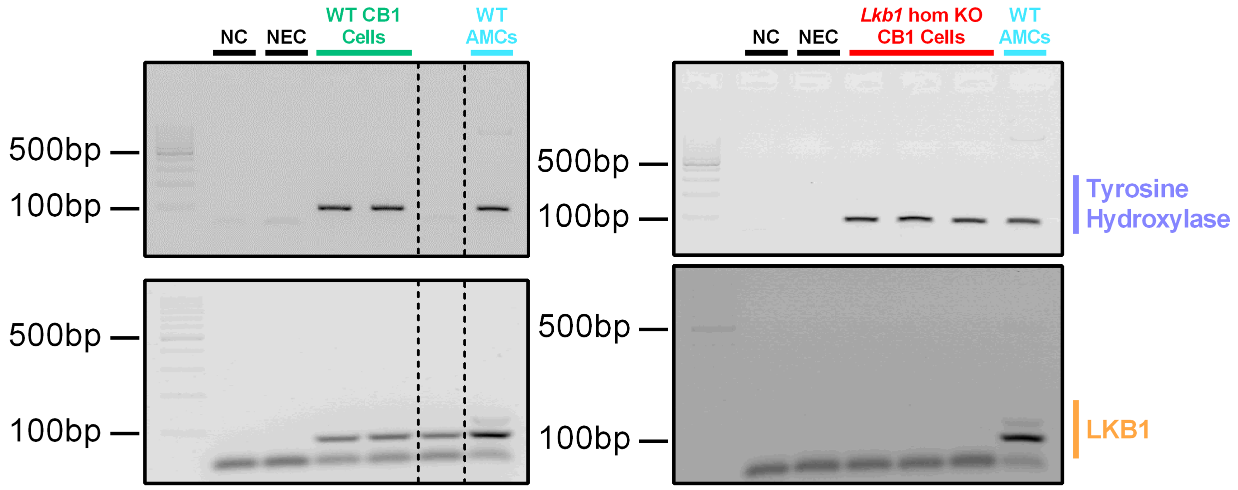


**Supplementary Figure 1.** Images show original gels for single cell end-point RT-PCR amplicons for tyrosine hydroxylase and LKB1 from acutely isolated adrenal medullary chromaffin cells from wild type mice (WT AMCs) and carotid body type I cells of wild type (WT CB1 cells**)** and conditional *Lkb1* knockout mice (*Lkb1* hom KO CB1 cells**)**; NC = negative control (cell aspirant but no reverse transcriptase added); NEC = negative extracellular control (aspirant of extracellular medium). This is one of three independent experiments.

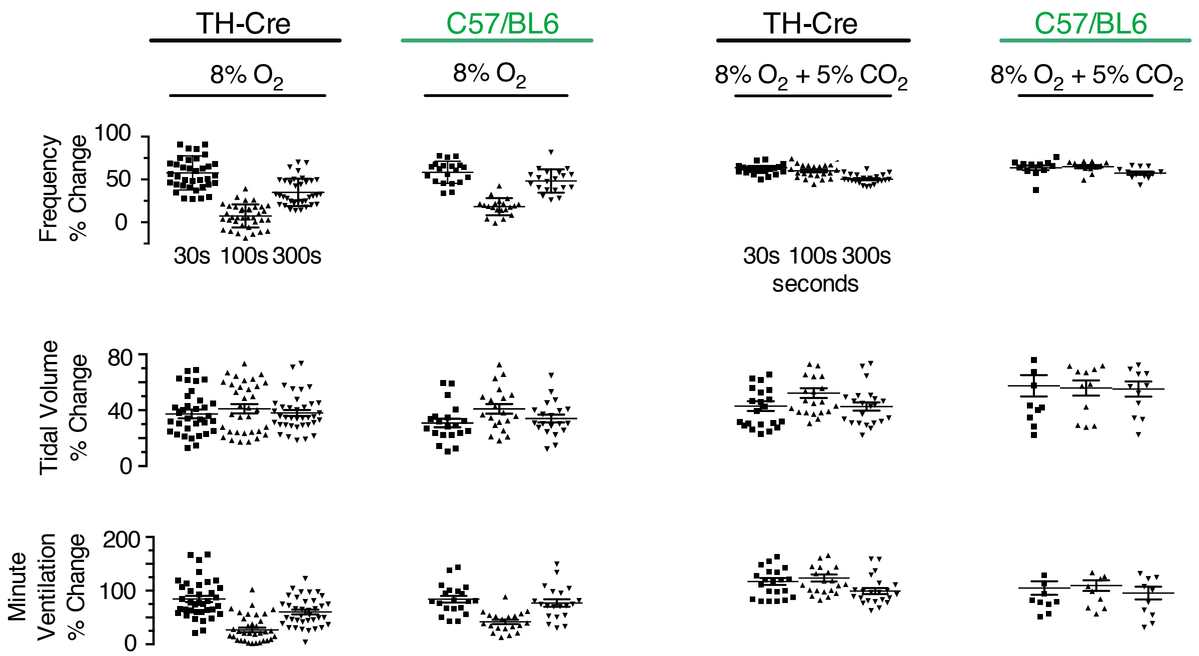


**Supplementary Figure 2. The mean ventilatory response to hypoxia is comparable between TH-Cre and C57/Bl6 mice.** Dot plots of mean±SEM for increases in minute ventilation (ml min$^{-1}$) at the peak of the Augmenting Phase (~30s), after Roll Off (~100s) and during the plateau of the Sustained Phase (~300s) of the ventilatory response to 12% and 8% $O_2$ for TH-Cre (black; 8% $O_2$, n = 25 independent experiments; $O_2$ + 5% $CO_2$, n = 17 independent experiments) and wild type C57/BL6 mice (green; 8% $O_2$, n = 20 independent experiments; 8% $O_2$ + 5% $CO_2$; n = 10 independent experiments).

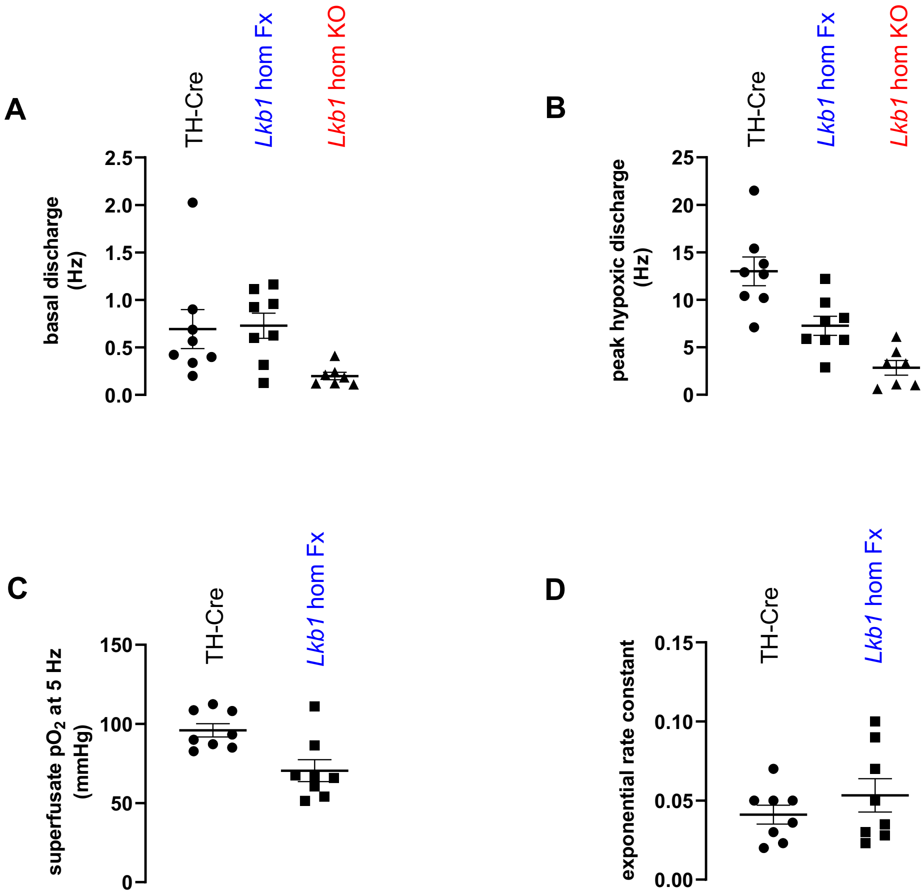


**Supplementary Figure 3. Conditional deletion of *Lkb1* in carotid body type I cells attenuates hypoxia-evoked afferent discharge from the carotid body in-vitro** A, Individual values and mean±SEM of basal/normoxic chemoafferent activity measured in TH-Cre (n = 8 different carotid bodies), homozygous *Lkb1* floxed (*Lkb1* hom Fx; n = 8 different carotid bodies) and conditional homozygous *Lkb1* knockout (*Lkb1* hom KO; n = 7 different carotid bodies) groups. B, Individual values and mean±SEM of peak hypoxic frequency. C, Individual values and mean±SEM of $PO_2$ required to achieve a frequency of 5 Hz, in TH-Cre and *Lkb1* hom Fx. D, Individual values and mean±SEM of hypoxic response curve exponential rate constants, in TH-Cre and *Lkb1* hom Fx groups.

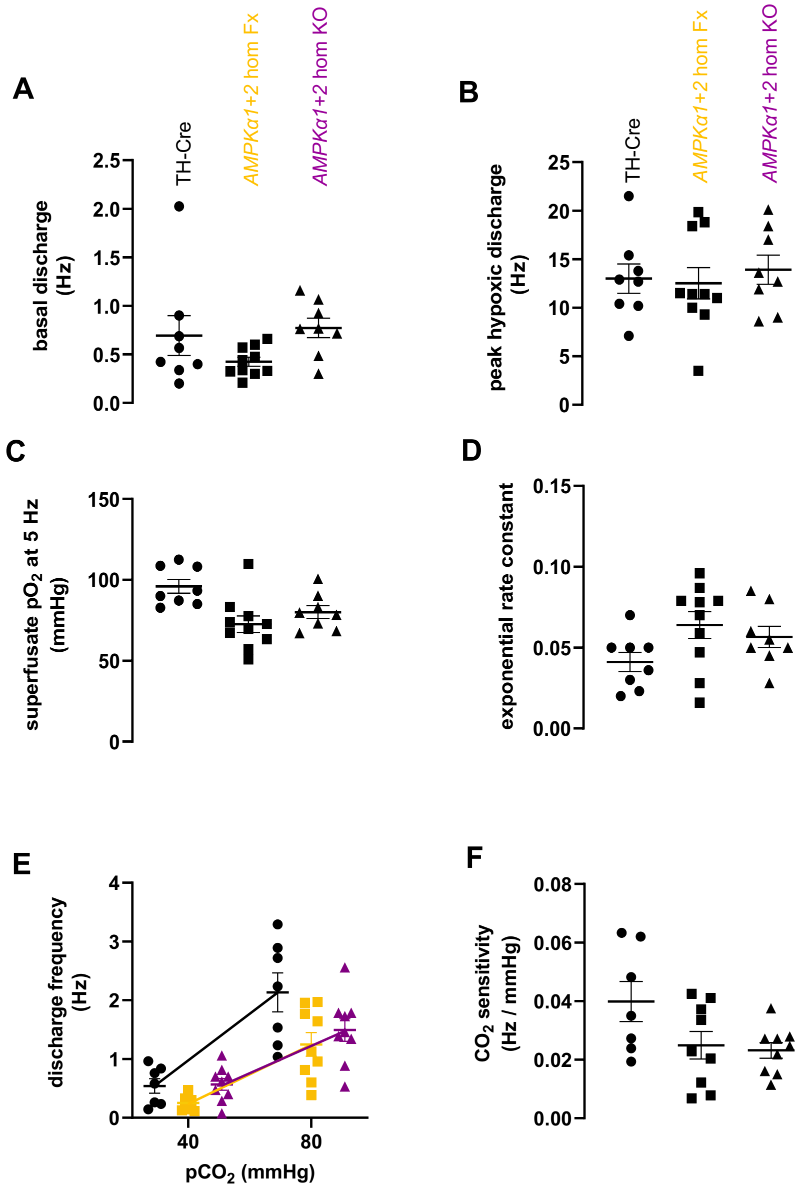


**Supplementary Figure 4. Conditional deletion of *AMPK-α1*+α2 in carotid body type I cells has no effect on hypoxia-evoked or hypercapnia evoked afferent discharge from the carotid body in-vitro** A-D, Individual values and mean±SEM of basal chemoafferent activity (A), peak hypoxic frequency (B), $PO_2$ at 5 Hz (C) and exponential rate constant (D) measured in TH-Cre (n = 8 different carotid bodies), *AMPKα1+α2* floxed *(AMPKα1+α2* hom Fx, n=10 different carotid bodies) and conditional homozygous *AMPKα1+α2* knockouts (*AMPKα1+α2* hom KO; n=8 different carotid bodies). E-F, Chemoafferent activity measured at a superfusate *P*$CO_2$ of 40 mmHg and 80 mmHg (hypercapnia) (E) and $CO_2$ sensitivity (F) for TH-Cre (n = 7 different carotid bodies), *AMPKα1+α2* hom Fx (n = 9 different carotid bodies) and *AMPKα1+α2* hom KO (n = 9 different carotid bodies) groups. The *P*$O_2$ required to reach a frequency of 5Hz was reduced in the *AMPKα1+α2* hom Fx (73±5 mmHg, $p<0.01$) and showed a similar trend in the *AMPKα1+α2* hom KO group (80±5 mmHg, $p = 0.08$) compared to TH-Cre controls (96±4 mmHg). There was, however, no difference between *AMPKα1+α2* hom Fx and *AMPKα1+α2* hom KO groups

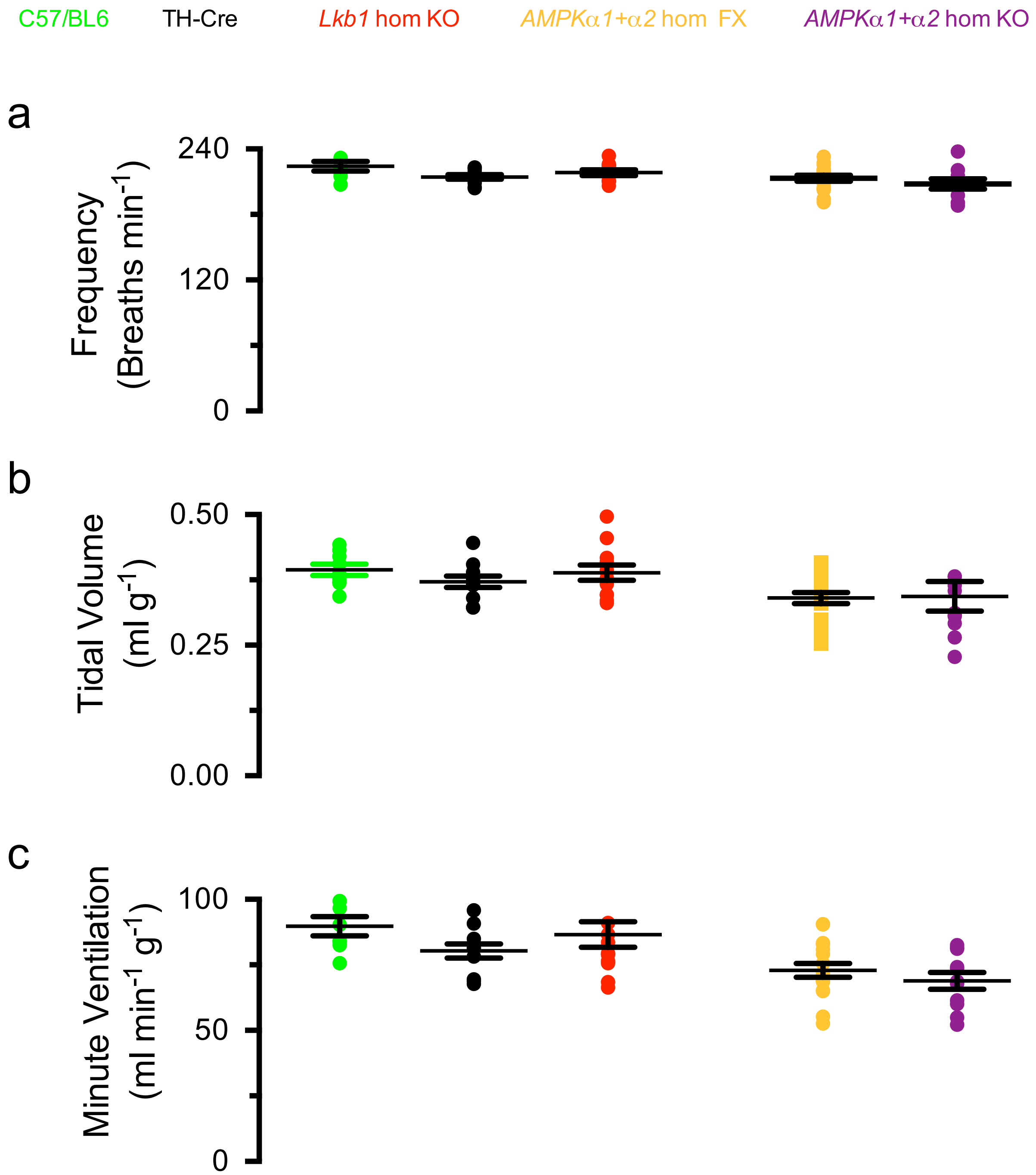


**Supplementary Figure 5. Basal breathing frequency, tidal volume and minute ventilation of experimental mice.** Bar charts show mean ± SEM for (A) breathing frequency ($min^{-1}$) (B) tidal volume ($ml\ g^{-1}$) and (C) minute ventilation ($ml\ min^{-1}\ g^{-1}$) under normoxia for C57/BL6 (green, n = 9 independent experiments), TH-Cre (black, n = 11 independent experiments), *Lkb1* homozygous floxed, *Lkb1* homozygous knockout (*Lkb1* hom KO, red; n = 12 independent experiments), *AMPKα1+α2* double Fx mice (*AMPKα1+α2* hom Fx; n = 17 independent experiments), *AMPKα1+α2* double knockout mice (*AMPKα1+α2* hom KO; n = 11 independent experiments).

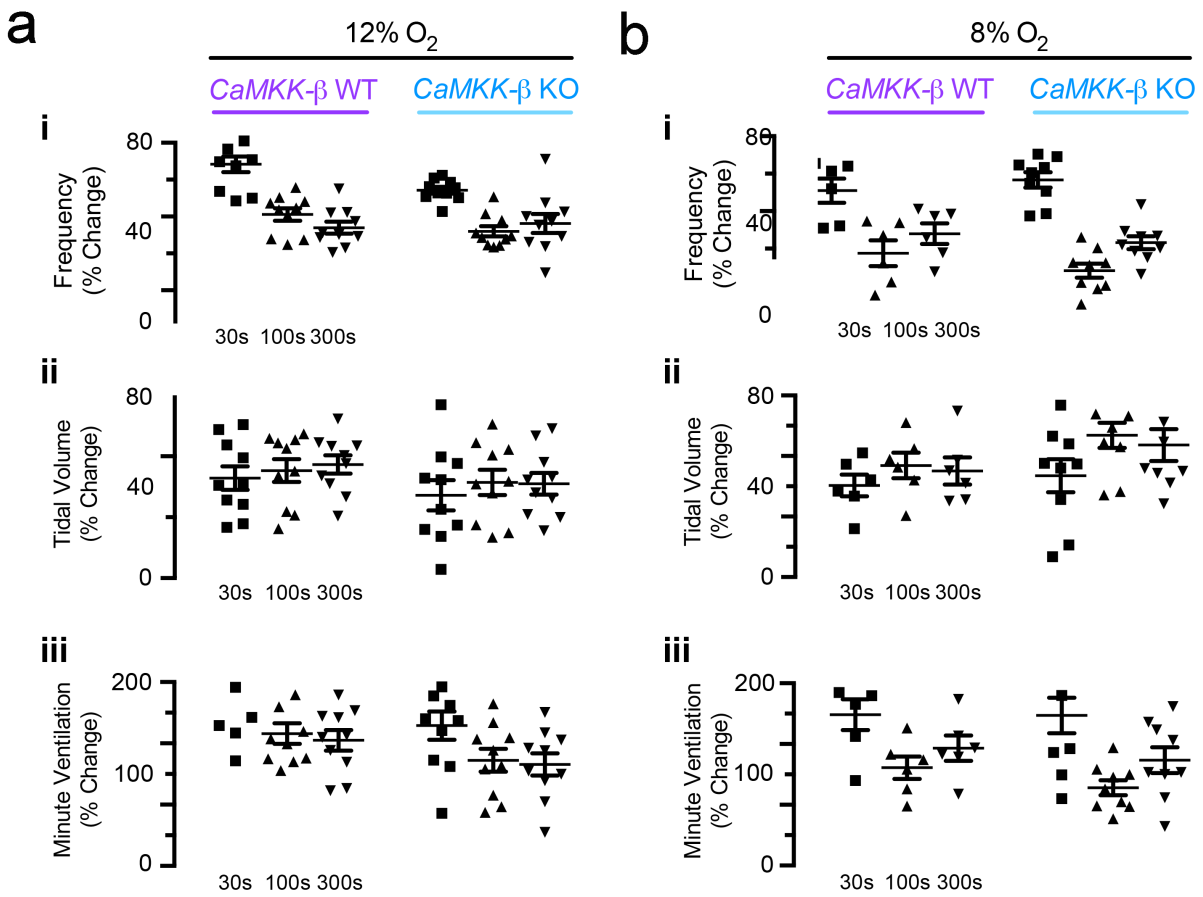


**Supplementary Figure 6. The hypoxic ventilatory response remains unaffected after global CaMKK2 deletion.** Dot plots and mean±SEM for *CaMKK2 WT* (n = 10 independent experiments) and global *CaMKK2* (CaMKK-β) KO mice (n = 10 independent carotid bodies) show increases in minute ventilation (ml $min^{-1}$) at the peak of the Augmenting Phase (A, ~30s), after Roll Off (RO, ~100s) and during the plateau of the Sustained Phase (SP, ~300s) of the ventilatory response to 12% and 8% $O_2$

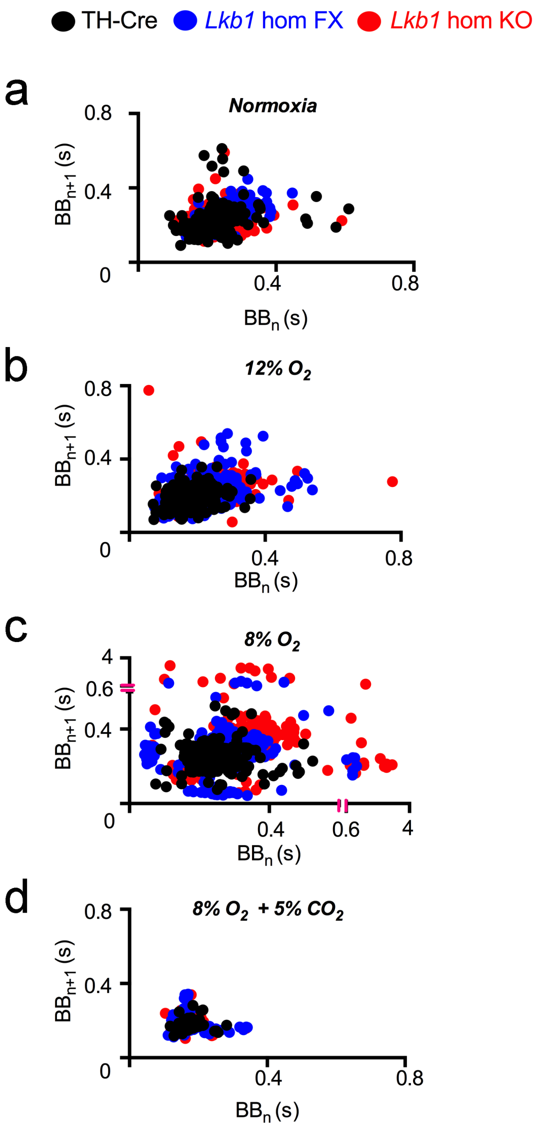


**Supplementary Figure 7. Respiratory dysfunction and apnoea are less severe for mice 90% hypomorphic for LKB1 than for conditional *Lkb1* knockouts**

a-d, Exemplar Poincaré plots of the inter-breath interval ($BB_n$) versus subsequent interval ($BB_{n+1}$) during (a) normoxia, (b) mild hypoxia (12% $O_2$), (c) severe hypoxia (8% $O_2$) and (d) severe hypoxic hypercapnia (8% $O_2$ + 5% $CO_2$;) for controls (TH-Cre, black; 12% $O_2$, n = 19 independent experiments; 8% $O_2$, n = 24 independent experiments; 8% $O_2$ + 5% $CO_2$, n = 17 independent experiments), *Lkb1 homozygous* floxed (*Lkb1* hom Fx, blue; n = 16 independent carotid bodies; 8% $O_2$, n = 15 independent experiments; 8% $O_2$ + 5% $CO_2$, n = 16 independent carotid bodies) and conditional *Lkb1* homozygous knockouts (*Lkb1* hom KO, red; 12% $O_2$, n = 17 independent experiments; 8% $O_2$, n = 28 independent experiments; 8% $O_2$ + 5% $CO_2$, n = 27 independent experiments).

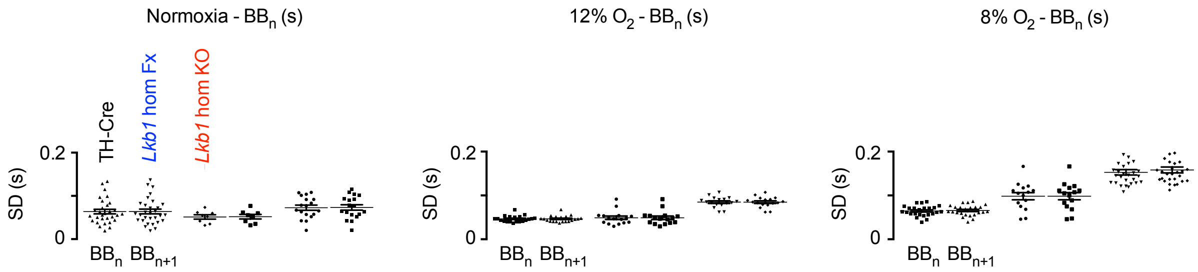


**Supplementary Figure 8. Respiratory dysfunction is less severe for mice that are ~90% hypomorphic for LKB1 than for conditional *Lkb1* knockouts.** Dot plots and mean±SEM for the standard deviation (SD) of $BB_n$ and $BB_{n+1}$ for controls (TH-Cre, black; 12% $O_2$, n = 19 independent experiments; 8% $O_2$, n = 24 independent experiments; 8% $O_2$ + 5% $CO_2$, n = 17 independent experiments), *Lkb1 homozygous* floxed (*Lkb1* hom Fx, blue; n = 16 independent carotid bodies; 8% $O_2$, n = 15 independent experiments; 8% $O_2$ + 5% $CO_2$, n = 16 independent carotid bodies) and conditional *Lkb1* homozygous knockouts (*Lkb1* hom KO, red; 12% $O_2$, n = 17 independent experiments; 8% $O_2$, n = 28 independent experiments; 8% $O_2$ + 5% $CO_2$, n = 27 independent experiments).

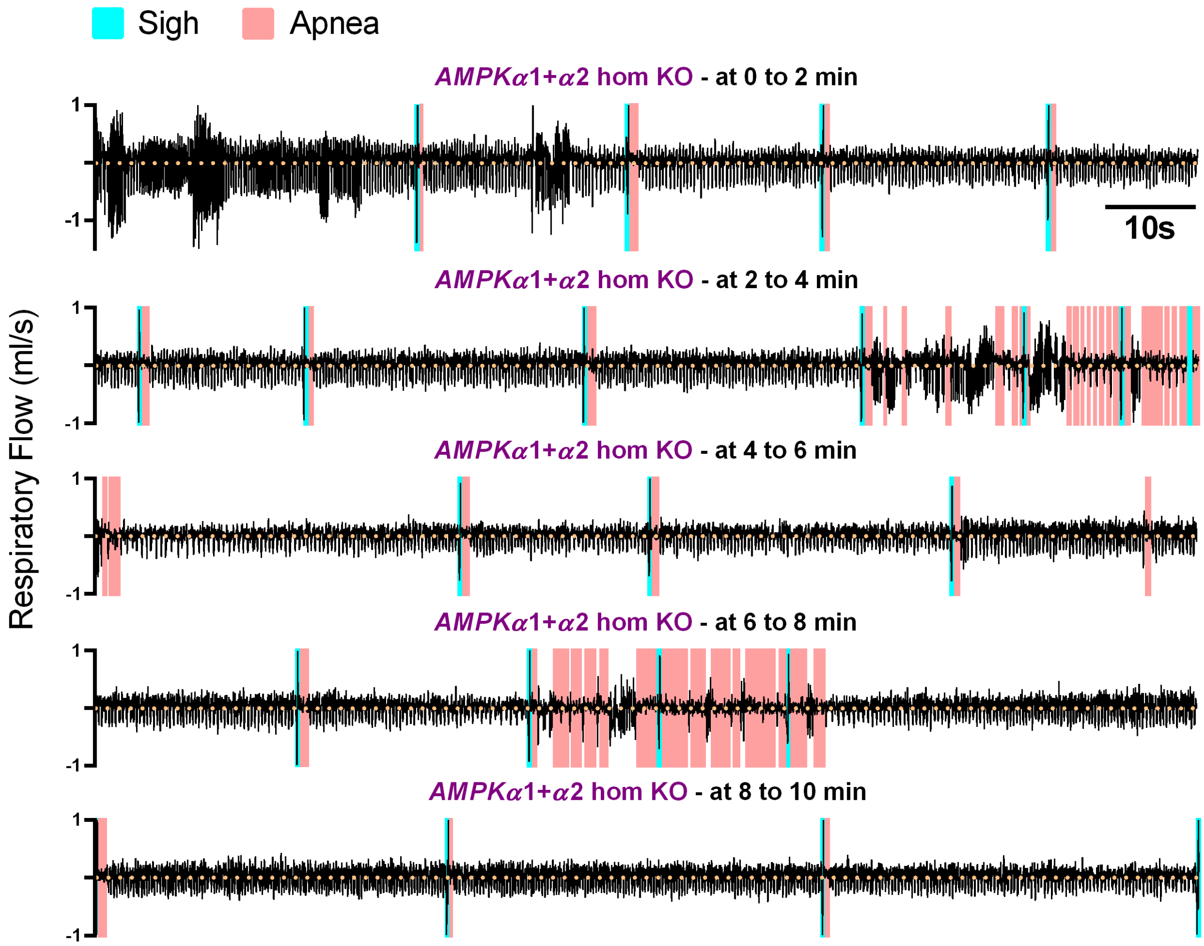


**Supplementary Figure 9 - Conditional deletion of *AMPKα1+α2* in tyrosine hydroxylase expressing cells does not precipitate Cheyne-Stokes-like ventilation even during prolonged exposure to severe hypoxia.**
Typical ventilatory record for conditional *AMPKα1+α2* double knockout mice (*AMPKα1+α2* hom KO) during a 10 minute exposure to severe hypoxia (8% $O_2$).

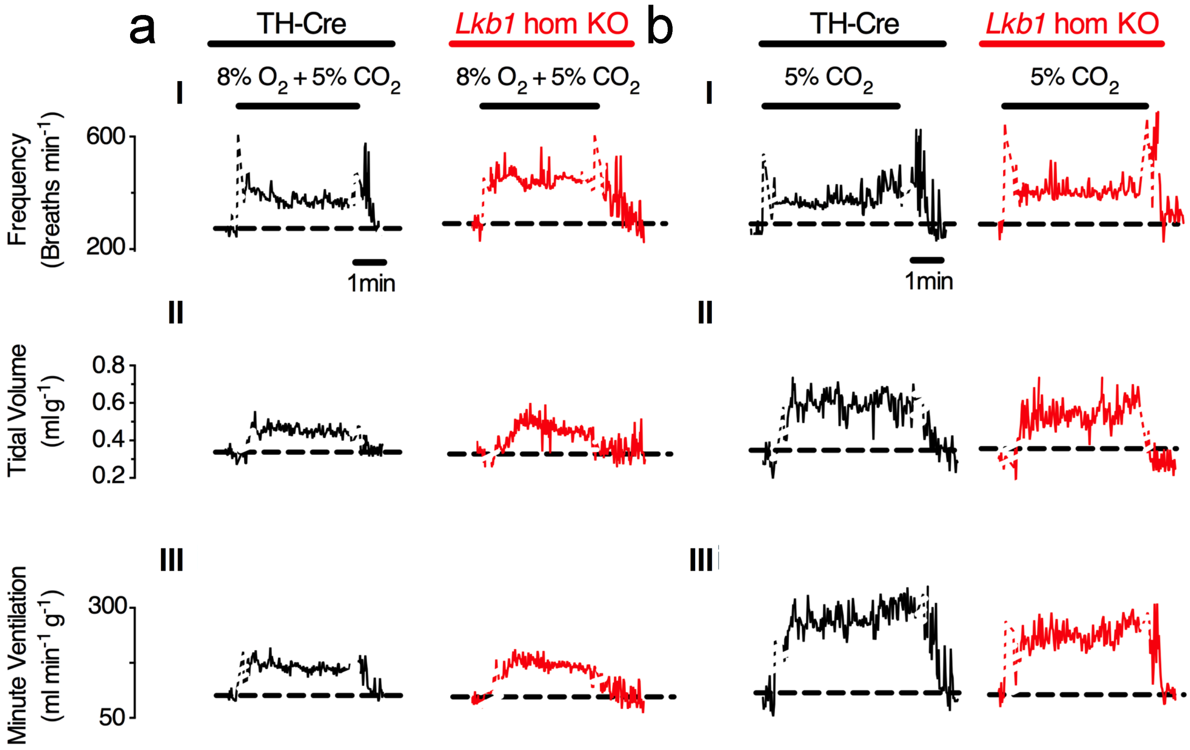


**Supplementary Figure 10 - *Lkb1* deletion attenuates the hypoxic ventilatory response in a manner that is $pO_2$-dependent but does not block the hypoxic-hypercapnic ventilatory responses or hypercapnic ventilatory responses.** Exemplar records show the responses over a 5min period of exposures to (A) severe hypoxia (8% $O_2$), (B) hypoxic hypercapnia (8% $O_2$ + 5% $CO_2$) and (C) hypercapnia (5% $CO_2$) on (I) breathing frequency ($min^{-1}$), (II) tidal volume (ml $g^{-1}$) and (III) minute ventilation (ml $min^{-1}$ $g^{-1}$) in TH-Cre (black) and *Lkb1* homozygous knockout mice (*Lkb1* hom KO, red) with 2s sampling periods. Dashed black line indicating basal level, white line breaks indicate artefact of gas exchange.

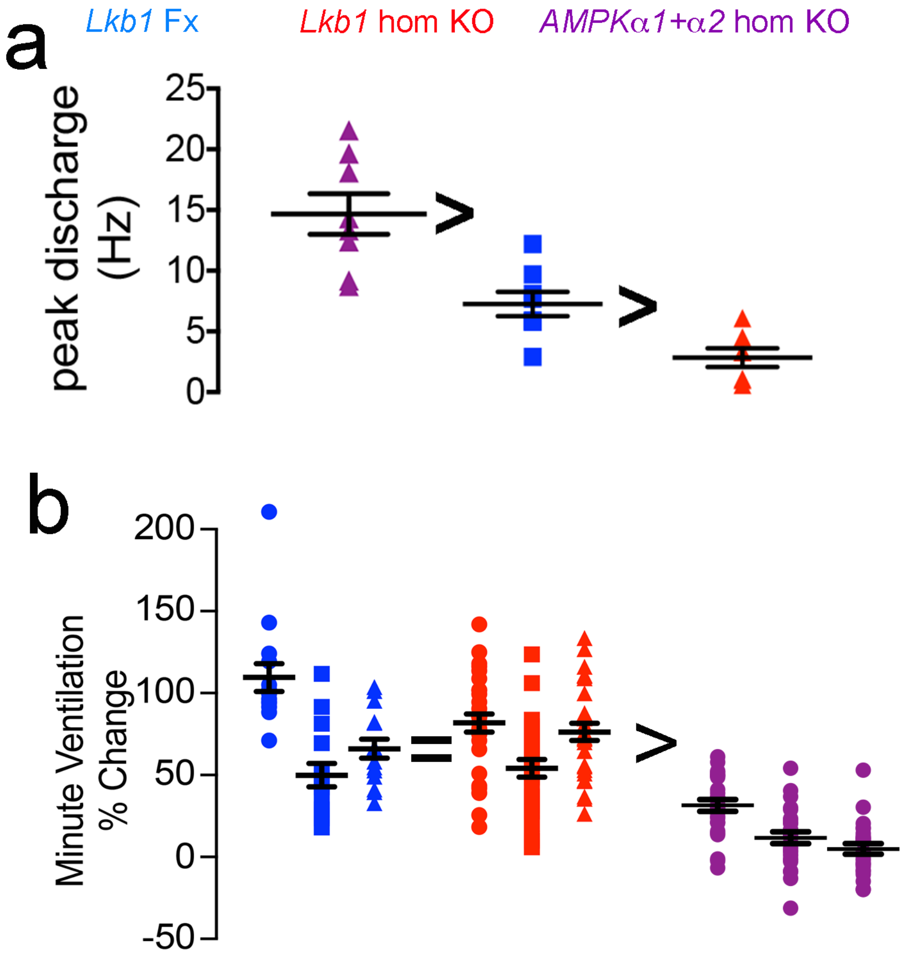


**Supplementary Figure 11 - Rank order of block of carotid body afferent fibre discharge and hypoxic ventilatory responses for LKB1 hypomorphs, *Lkb1* and *AMPKα1*+α2 knockouts.**

a, Comparison of mean±SEM for peak carotid body afferent fibre discharge for *Lkb1* homozygous floxed mice (blue, n = 8 different carotid bodies), *Lkb1* homozygous knockouts (red, n = 7 different carotid bodies), and conditional *AMPKα1*+α2 knockouts (mauve, n = 9 different carotid bodies). *=p<0.05, **=p<0.01, ***p=<0.001, ****=p<0.0001. b, Comparison of mean±SEM for the % change in minute ventilation at the peak of the Augmenting Phase (AP, ~30s), at ~100s following Roll Off (RO) and during the plateau of the Sustained Phase (SP, ~300s) during exposures to 8% $O_2$ of *Lkb1* homozygous floxed mice (*Lkb1* hom Fx, blue; n = 15 independent experiments), conditional *Lkb1* knockouts (*Lkb1* hom KO, red; n = 30 independent experiments), and *AMPKα1+α2* knockouts (*AMPKα1+α2* hom KO, purple; n = 26 independent experiments).

| **Table 1 Core body temperature, venous blood gas, and blood pH analysis** | | | | | | | | |
|---|---|---|---|---|---|---|---|---|
| | C57/Bl6 | TH-Cre | *Lkb1* hom KO | *AMPK* Double Fx | *AMPK α1* KO | *AMPK α2* KO | *AMPK* Double KO | P value |
| Temperature (°C) | 34.4±0.3 | 34.9±0.5 | 34.7±0.3 | 35.8±0.4 | 35.6±0.4 | 36.4±0.4 | 35.6±0.4 | NS |
| $PO_2$ (mmHg) | 43.7±8.9 | 60.7±10.5 | 34.7±2.6 | 52.2±2.5 | 53.8±9.4 | 47.0±6.8 | 57.3±8.9 | NS |
| $PCO_2$ (mmHg) | 64.2±3.6 | 80.7±7.0 | 71.2±7.1 | 52.5±5.5 | 54.3±10.0 | 57.5±2.1 | 64.8±6.7 | NS |
| pH | 7.1±0.03 n=4 | 7.1±0.03 n=3 | 7.1±0.03 n=3 | 7.2±0.02 n=5 | 7.2±0.09 n=4 | 7.2±0.01 n=4 | 7.1±0.02 n=4 | NS |

Values shown as mean±SEM
Significance tested by one-way ANOVA
